# EphB6 regulates social behavior through gut microbiota-mediated vitamin B6 metabolism and excitation/inhibition balance of medial prefrontal cortex in mice

**DOI:** 10.1101/787622

**Authors:** Ying Li, Zheng-Yi Luo, Yu-Ying Hu, Yue-Wei Bi, Jian-Ming Yang, Ming-An Liu, Wen-Jun Zou, Shi Li, Tong Shen, Shu-Ji Li, Lang Huang, Ai-Jun Zhou, Yun-Long Song, Tian-Ming Gao, Jian-Ming Li

**Author notes:** These authors contributed equally to this work (Ying Li, Zheng-Yi Luo). Correspondence to: Jian-Ming Li, Department of Pathology, Sun Yat-Sen Memorial Hospital, Guangzhou 510120, People’s Republic of China. Or Tian-Ming Gao, State Key Laboratory of Organ Failure Research, Key Laboratory of Psychiatric Disorders of Guangdong Province, Department of Neurobiology, School of Basic Medical Sciences, Southern Medical University, Guangzhou 510515, People’s Republic of China.

## Abstract

Autism spectrum disorder (ASD) is a developmental disorder with no effective pharmacological treatments so far. Gut microbiota has been suggested to contribute to autistic symptoms. However, the key genes and the mechanisms linking gut microbiota and brain dysfunctions in ASD are still unclear. Here, we found deletion of EphB6, an ASD-associated candidate gene, induced dysregulated gut microbiota and autism-like behavior in mice. More importantly, transplanting fecal microbiota from EphB6-deficient mice resulted in disturbed gut microbiota and autism-like behavior in antibiotics-treated C57BL/6J mice. Meanwhile, transplanting fecal microbiota from wild-type mice ameliorated disturbed gut microbiota and autism-like behavior in mice with deletion of EphB6. At the metabolic levels, dysregulated gut microbiota led to vitamin B6 and dopamine defects in EphB6-deficient mice. At the cellular levels, excitation/inhibition (E/I) imbalance in medial prefrontal cortex was induced by gut microbiota-mediated defects of vitamin B6 metabolism in EphB6-deficient mice. Our study uncovers a key role for EphB6 in regulation of social behavior by gut microbiota-mediated vitamin B6 metabolism, dopamine synthesis and E/I balance, suggesting a new strategy for treatment of ASD patients.

## Introduction

Autism spectrum disorder (ASD), affecting approximately 1% of the population around the world, is mainly characterized with impaired social interaction and communication, and restricted and repetitive behavior (Lord, Elsabbagh, Baird, & Veenstra-Vanderweele, 2018). So far, only early behavioral and educational interferences show ameliorative effects on autistic symptoms of ASD patients and there are no effective pharmacological therapies for the treatment of core autistic symptoms (Lord et al., 2018).

It is generally considered that ASD is caused by genetic, developmental and environmental factors. And the genetic factor is considered to be the most important cause of ASD. With the development of sequencing technique, more and more genes have been found to be related with ASD.

EphB6, belonging to the Eph family of receptor tyrosine kinases, plays an important role in regulating Eph receptor signaling networks, T cell functions, development of intestinal epithelium and epithelial homeostasis (Luo, Yu, Tremblay, & Wu, 2004; Miao & Wang, 2009; Truitt & Freywald, 2011). EphB6 has been suggested as a candidate of ASD-associated-gene (Butler, Rafi, & Manzardo, 2015; Chen, 2009) and a recent genomic study has found the mutation of EphB6 in some ASD patients (O’Roak et al., 2011). Nevertheless, the role of EphB6 and the mechanisms involved in ASD are still unclear.

Accumulating evidence shows that gut-brain-microbiota axis plays a key role in regulating homeostasis of human body. Gut microorganisms have been reported to participate in a lot of neuropsychiatric disorders, such as anxiety disorders, depression (Foster & McVey Neufeld, 2013) and epilepsy (Olson et al., 2018). In most ASD patients, changed gut microorganisms and serious gastrointestinal problems are observed (Coury et al., 2012; Parracho, Bingham, Gibson, & McCartney, 2005; Sharon et al., 2019). Interestingly, several studies have found the important role of gut microbiota in modulating ASD-like phenotypes of mice (Buffington et al., 2016; Hsiao et al., 2013; Sgritta et al., 2019). A clinical study shows that microbiota transfer therapy can improve gastrointestinal problems and autistic symptoms of ASD patients, who are 7 to 16 years old, and this benefit can last for two years (D.-W. Kang et al., 2017; D. W. Kang et al., 2019). These studies suggest that gut-brain-microbiota axis may have a significant impact on the development of ASD. However, how gut microbiota contributes to dysregulation of brain function is still not well-characterized.

In our study, we find EphB6 is functionally associated with ASD and regulates social behavior by gut microbiota-mediated vitamin B6 and dopamine metabolism. More importantly, we establish the functional link between dysregulated gut microbiota and excitation/inhibition (E/I) imbalance in medial prefrontal cortex (mPFC), a key gut-brain functional axis, in EphB6-deficient mice.

## Results

### Deletion of EphB6 led to autism-like behavior and gut microbial disturbance in mice

As a candidate gene associated with ASD, whether and how EphB6 works in ASD is still unclear. To answer these questions, we established EphB6 knockout mice. We found a deletion of EphB6 in different tissues, including colon, brain, lung and spleen, in EphB6 knockout mice (KO mice) compared with EphB6^+/+^ (wild-type, WT) mice (Supplementary Fig. 1C-D). However, the weight of brain and body, the length of body and daily dietary consumption were similar between the two groups of mice in spite of the deletion of EphB6 (Supplementary Fig. 1E-H).

Patients with ASD often display repetitive stereotyped behavior and social deficits. Interestingly, we found KO mice spent more time on self-grooming compared with WT mice (Fig. 1A). While in marble burying test, KO mice buried similar marbles with WT mice (Supplementary Fig. 1J). In social partition test, KO mice spent less time on sniffing at partition, no matter the familiar or novel mouse was put in, than WT mice (Fig. 1B). In three-chambered social approach task, KO mice spent similar time in chambers with an unfamiliar mouse or inanimate object (Fig. 1D), also KO mice showed less preference for the social mouse (stranger 1) over the object than the WT mice (Fig. 1F-G). While the novel social partner (stranger 2) was put into the empty wire cage, KO mice still spent similar time in the two chambers (Fig. 1E) and showed less preference for the novel mouse over the familiar mouse than WT mice (Fig. 1H). These results confirmed the abnormalities of social interaction in KO mice sufficiently. Olfactory cues have been generally considered to be of the most importance in communication among mice (Carter, Williams, Witt, & Insel, 1992; Keverne, 2002). In olfactory habituation/dishabituation test, repeated presentation of cotton swabs saturated by same odor caused less and less time spent sniffing at cotton swabs and presentation of cotton swab saturated by a new odor caused increased time spent sniffing both in WT and KO mice. However, when introducing a cotton swab saturated with social odor, KO mice showed less interest in social odors than WT mice (Fig. 1I). These results implied the communication deficits in KO mice although the ability to discriminate and habituate different odors was normal.

**Fig. 1.**
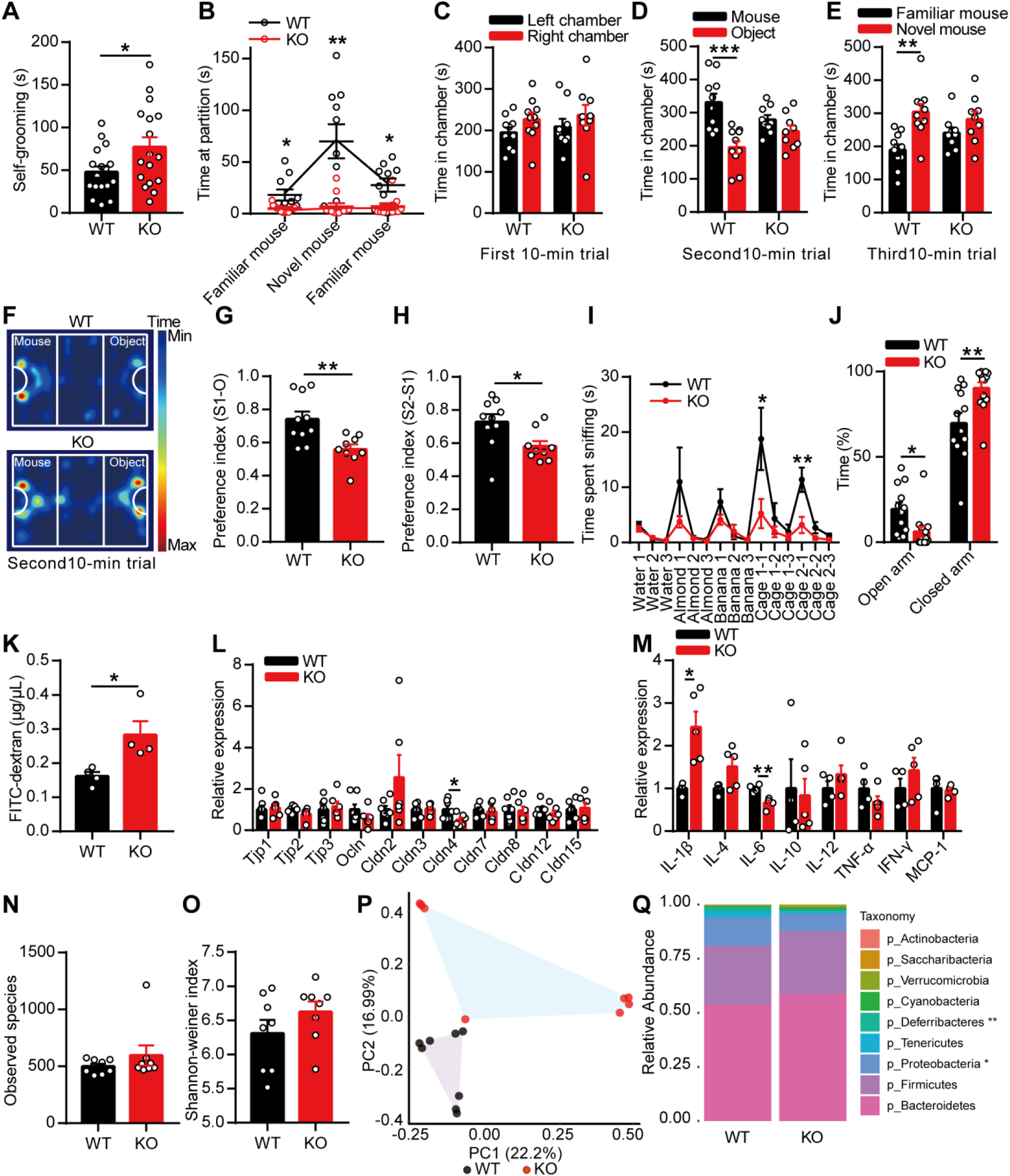
Deletion of EphB6 led to autism-like behavior and gut microbial disturbance in mice. (A) 8-week-old male KO mice spent more time on self-grooming compared with WT mice. n = 17 mice for each group. (B) In social partition test, KO mice spent less time on sniffing mice compared with WT mice. n = 10, 11 mice respectively. (C-H) In three-chambered social approach task, time spent in chambers during different 10-min trials (C-E), trajectory diagram during the second 10-min trial (F) were showed. KO mice showed less preference for the social mouse over the object (G) and less preference for the novel social mouse over the familiar social mouse (H) compared with WT mice. n = 10, 9 mice respectively. (I) In olfactory habituation/dishabituation test, KO mice spent less time on sniffing social odors compared with WT mice. n = 11 mice for each group. (J) In elevated-plus-maze test, KO mice spent less time in open arm and more time in closed arm compared with WT mice. n = 12, 13 mice respectively. (K) The intestinal permeability of 8-week-old WT and KO mice was detected using FITC-dextran. n = 4 mice for each group. (L) The mRNA expressions of tight junction molecules were detected in colon of 8-week-old WT and KO mice. n = 7, 6 mice respectively. (M) The mRNA expressions of cytokines were detected in colon of 8-week-old WT and KO mice. n = 4, 5 mice respectively. (N-Q) 16S rDNA gene sequencing of gut microbiota of 8-week-old WT and KO mice. The species richness (N) and diversity (O) of gut microbiota were similar, while the microbial composition (P) was different between the two groups. Relative abundance of different bacteria in phylum level was showed (Q). n = 8 mice for each group. Data shown are mean ± SEM. Two-tailed unpaired student’s *t* test (A, C-E, G-H, J-O), two-way repeated measures ANOVA (B, I), anosim analysis (P). *, p < 0.05; **, p < 0.01; ***, p < 0.001. WT, EphB6^+/+^ mice; KO, EphB6^−/−^ mice; GI, gastrointestinal tract; FITC, fluorescein isothiocyanate. Statistical values are presented in Supplementary Table 2.

Patients with ASD are often accompanied with other mental diseases, such as hyperactivity, anxiety and intellectual disability. In open field test, KO mice showed same locomotor activities and spent almost same time in center area compared with WT mice (Supplementary Fig. 1K-L). While in elevated-plus-maze test, KO mice spent less time in open arm and more time in closed arm compared with WT mice (Fig. 1J), which implied that KO mice displayed anxiety-like behavior. In morris water maze, KO mice had the normal spatial learning and memory as the WT mice (Supplementary Fig. 1M-O). Collectively, mice with deletion of EphB6 showed autism-like behavior, including stereotyped behavior and social deficits, accompanied with anxiety-like behavior.

Eph/ephrin signaling has been reported to modulate gut epithelial development and homeostasis. Also, it has been generally accepted that many ASD patients have gastrointestinal (GI) symptoms (Buie et al., 2010; Coury et al., 2012; Parracho et al., 2005) and a changed composition of gut microbiota (Sharon et al., 2019). Then we want to know whether mice with deletion of EphB6 will suffer from GI problems. To measure intestinal permeability by FITC-dextran, we found the significantly increased intestinal permeability in EphB6-deficient mice compared with WT mice (Fig. 1K). Accordingly, the mRNA expression of Cldn4, one member of tight junction molecules, was decreased in colon of KO mice compared with WT mice (Fig. 1L). In addition, we detected that mRNA expression of IL-1β, as a proinflammatory factor, was dramatically increased and IL-6, which has anti-inflammatory effect, was decreased in colon of KO mice compared with WT mice (Fig. 1M). GI problems in KO mice was not accompanied with morphological changes in small intestine, colon, kidney, liver, thymus and lung (Supplementary Fig. 1P).

The integrity of intestinal mucosa was important to maintain the balance of ecological environment in animals’ gut. Then we detected the fecal microbial populations of mice using 16S rDNA gene sequencing. There were no differences in microbial species richness and diversity between the two groups (Fig. 1N-O). Notably, principle coordinates analysis of Bray-Curtis distance showed fecal microbiota of KO mice clustered differently from WT mice (Fig. 1P), which predicted a different gut microbial composition between the two groups. In phylum level, the differences between the two groups were caused by the decreased *Deferribacteres* and *Proteobacteria* in fecal microbiota of KO mice (Fig. 1Q). In general, our results indicated deletion of EphB6 resulted in increased intestinal permeability and changes of gut microbial composition in mice.

Many studies have referred that GI problems and abnormal behavior of ASD always appear parallelly in patients (Parracho et al., 2005). Then we wondered when the gut microbial composition began to change in KO mice. We found the microbial species richness and diversity were also similar between 3-week-old and 4-week-old WT and KO mice (Supplementary Fig. 2A-B). In principle coordinates analysis, gut microbiota of 4-week-old KO mice clustered differently from 4-week-old WT mice (Supplementary Fig. 2D), while gut microbiota of 3-week-old WT and KO mice clustered similarly (Supplementary Fig. 2C). At the same time, 4-week-old KO mice, but not 3-week-old KO mice, showed increased self-grooming and decreased interest in social odor compared with even-aged WT mice (Supplementary Fig. 2E-G). These results further implied the possible relation between the abnormal behavior and gut microbial dysbiosis in mice with deletion of EphB6.

### Transplantation of fecal microbiota from EphB6-deficient mice caused autism-like behavior in SPF C57BL/6J mice

ASD is generally considered to be a neuro-developmental disorder, postnatal developmental disorder can also cause autism in patients (Zoghbi, 2003), and postnatal mutation of Nrxn1 in neurons led to autism-like behavior in mice (Rabaneda, Robles-Lanuza, Nieto-Gonzalez, & Scholl, 2014). Also gut microbiota of ASD patients could induce autism-like behavior in mice (Sharon et al., 2019). So, to study the relation between gut microbial dysbiosis and autism-like behavior in mice with deletion of EphB6, we gavaged the fecal microbiota from 8-week-old male WT or KO mice to 3-week-old SPF male C57BL/6J mice for a week (Fig. 2A). Three weeks after the gavage of fecal microbiota, gut microbial composition in SPF C57BL/6J mice treated with fecal microbiota from WT and KO mice was different (Fig. 2B-D). More interestingly, C57BL/6J mice with the gastric perfusion of fecal microbiota from KO mice displayed increased self-grooming (Fig. 2E) and decreased social behavior (Fig. 2F-H) compared with control mice. While in open field test and elevated-plus-maze test, the two groups of mice behaved similarly (Supplementary Fig. 2H-J). Furthermore, we gavaged orally the suspending solution of fecal microbiota from WT or KO mice to antibiotic-pretreated SPF male C57BL/6J mice. After pretreatment with antibiotics for 5 days, fecal microbiota of 8-week-old male WT or KO mice was gavaged orally to 3-week-old SPF male C57BL/6J mice for 5 days (Fig. 2J). About 2 weeks after fecal microbial colonization, similarly, we found gut microbiota of SPF C57BL/6J mice treated with fecal microbiota from KO mice clustered differently from control mice (Fig. 2K-M). Then we found C57BL/6J mice with gastric perfusion of fecal bacteria from KO mice showed increased self-grooming (Fig. 2N) and decreased social behavior (Fig. 2O-R). Also, the two groups of mice behaved similarly in open field test and elevated-plus-maze test (Supplementary Fig. 2K-M). What’s more, fecal microbiota from 4-week-old KO mice, but not 3-week-old, induced increased self-grooming and social deficits in 3-week-old SPF C57BL/6J mice compared with C57BL/6J mice gavaged with fecal microbiota from even-aged WT mice (Supplementary Fig. 2N-S). Collectively, fecal microbiota from EphB6-deficient mice caused more self-grooming and impaired social behavior in C57BL/6J mice.

**Fig. 2.**
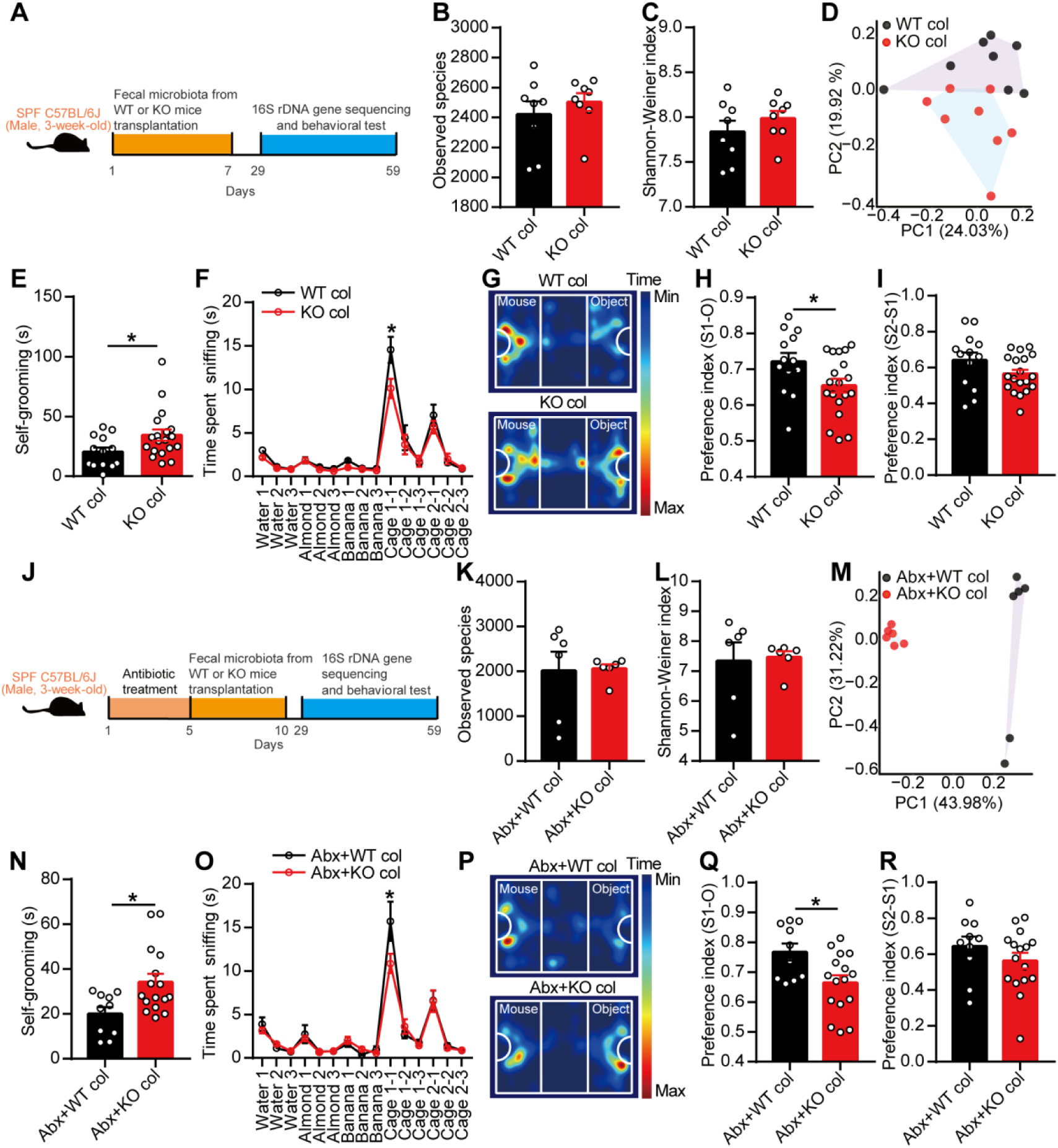
Transplantation of fecal microbiota from EphB6-deficient mice caused autism-like behavior in SPF C57BL/6J mice. (A-I) Schematic of the fecal microbiota transplantation (A). 3-week-old SPF male C57BL/6J mice were gavaged orally with fecal microbiota from 8-week-old male WT or KO mice (each contained 8 healthy mice from at least 3 cages) for a week. After 3 weeks, fecal microbiota of the treated C57BL/6J mice were sequenced (B-D, 8 treated C57BL/6J mice of each group were selected randomly from at least 3 cages) and self-grooming test (E), olfactory habituation/dishabituation test (F), three-chambered social approach task (G-I), open field test and elevated-plus-maze test were conducted with an interval of at least 2 days (E-I, n = 13, 19 mice respectively). (J-R) Schematic of fecal microbiota transplantation (J). 3-week-old SPF male C57BL/6J mice were gavaged orally with antibiotics (ampicillin, vancomycin, neomycin, metronidazole) for 5 days and then gavaged orally with fecal microbiota from 8-week-old male WT or KO mice (each contained 8 healthy mice from at least 3 cages) for another 5 days. After 19 days, fecal microbiota of the treated C57BL/6J mice were sequenced (K-M, 6 treated C57BL/6J mice of each group were selected randomly from at least 2 cages) and self-grooming test (N), olfactory habituation/dishabituation test (O), three-chambered social approach task (P-R), open field test and elevated-plus-maze test were conducted with an interval of at least 2 days (N-R, n = 10, 16 mice respectively). Data shown are mean ± SEM. Two-tailed unpaired student’s *t* test (B-C, E, H-I, K-L, N, Q-R), two-way repeated measures ANOVA (F, O), anosim analysis (D, M). *, p < 0.05. WT col or KO col, colonized with fecal microbiota from EphB6^+/+^ or EphB6^−/−^ mice; Abx, pre-treated with antibiotics (ampicillin, vancomycin, neomycin, metronidazole). Statistical values are presented in Supplementary Table 2.

Then we wondered whether gut microbiota still played the role in autism-like behavior in adult mice. To begin, we gavaged orally a mixture of antibiotics to 6-week-old male SPF C57BL/6J mice for a week. And we found antibiotic treatment disrupted the gut microbiota greatly and induced decreased self-grooming and social deficits in adult C57BL/6J mice (Fig. 3A-I). These results indicated us that gut microbiota was related with autism-like behavior even in adult mice and different gut microbiota probably contributed to different behaviors, such as self-grooming and social behavior. Then, we gavaged the fecal microbiota from 8-week-old male WT or KO mice directly to 6-week-old SPF male C57BL/6J mice for a week. And we found fecal microbiota from KO mice also induced disturbed gut microbiota, more self-grooming and social deficits in adult C57BL/6J mice (Fig. 3J-R).

**Fig. 3.**
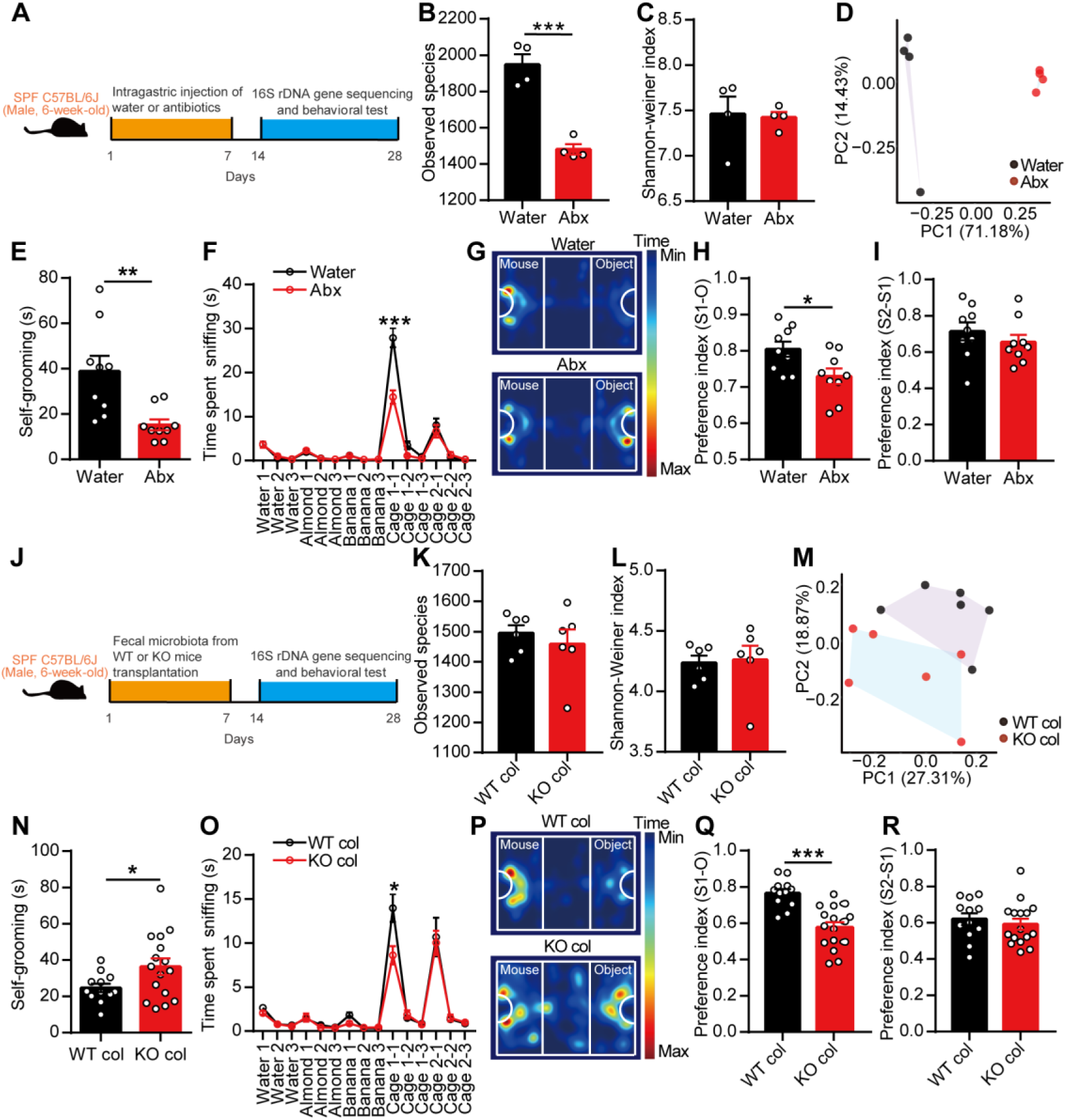
Antibiotic treatment and fecal microbiota transplantation from EphB6-deficient mice induced social deficits in 6-week-old SPF C57BL/6J mice. (A-I) Schematic of the treatment of antibiotics (A). 6-week-old SPF male C57BL/6J mice were gavaged orally with antibiotics (ampicillin, vancomycin, neomycin, metronidazole) for 7 days. After 7 days, the fecal microbiota of the treated C57BL/6J mice were sequenced (B-D, n = 4 mice for each group) and self-grooming test (E), olfactory habituation/dishabituation test (F), three-chambered social approach task (G-I) were conducted with an interval of at least 2 days. n = 9 mice for each group. (J-R) Schematic of the fecal microbiota transplantation (J). 6-week-old SPF male C57BL/6J mice were gavaged orally with fecal microbiota from 8-week-old male WT or KO mice for a week. After a week, the fecal microbiota of the treated C57BL/6J mice were sequenced (K-M, n = 6 mice for each group) and self-grooming test (N), olfactory habituation/dishabituation test (O), three-chambered social approach test (P-R) were conducted with an interval of at least 2 days (N-R, n = 12, 16 mice respectively). Data shown are mean ± SEM. Two-tailed unpaired student’s *t* test (B-C, E, H-I, K-L, N, Q-R), two-way repeated measures ANOVA (F, O), anosim analysis (D, M). *, p < 0.05; **, p < 0.01; ***, p < 0.001. Abx, treated with antibiotics (ampicillin, vancomycin, neomycin, metronidazole); WT col or KO col, colonized with fecal microbiota from EphB6^+/+^ mice or EphB6^−/−^ mice. Statistical values are presented in Supplementary Table 2.

Basically, our results indicated the important role of gut microbiota in autism-like behavior, even in adult mice.

### Transplantation of fecal microbiota from healthy wild-type mice successfully ameliorated autism-like behavior in adult mice with deletion of EphB6

Until now, there has no studies focusing on the effectiveness of microbiota transplantation in adult ASD patients. Then we gavaged orally the fecal microbiota from 8-week-old male WT mice to 8-week-old KO mice for a week. A week later, we found the gut microbiota of KO mice gavaged with fecal microbiota of WT mice clustered differently from KO mice gavaged with sterile PBS (Fig. 4B). In phylum level, we found the relative abundance of *Deferribacteres* was increased in KO mice gavaged with fecal microbiota of WT mice (Fig. 4C-D). And in species level, we found *Mucispirillum*, which is a genus in the phylum *Deferribacteres*, was increased in KO mice treated with fecal microbiota of WT mice (Fig. 4E). Also, fecal microbiota transplantation ameliorated the decreased relative abundance of *Prevotellaceae_UCG-001* and the increased relative abundance of *Lactobacillales* in KO mice (Fig. 4F-G).

**Fig. 4.**
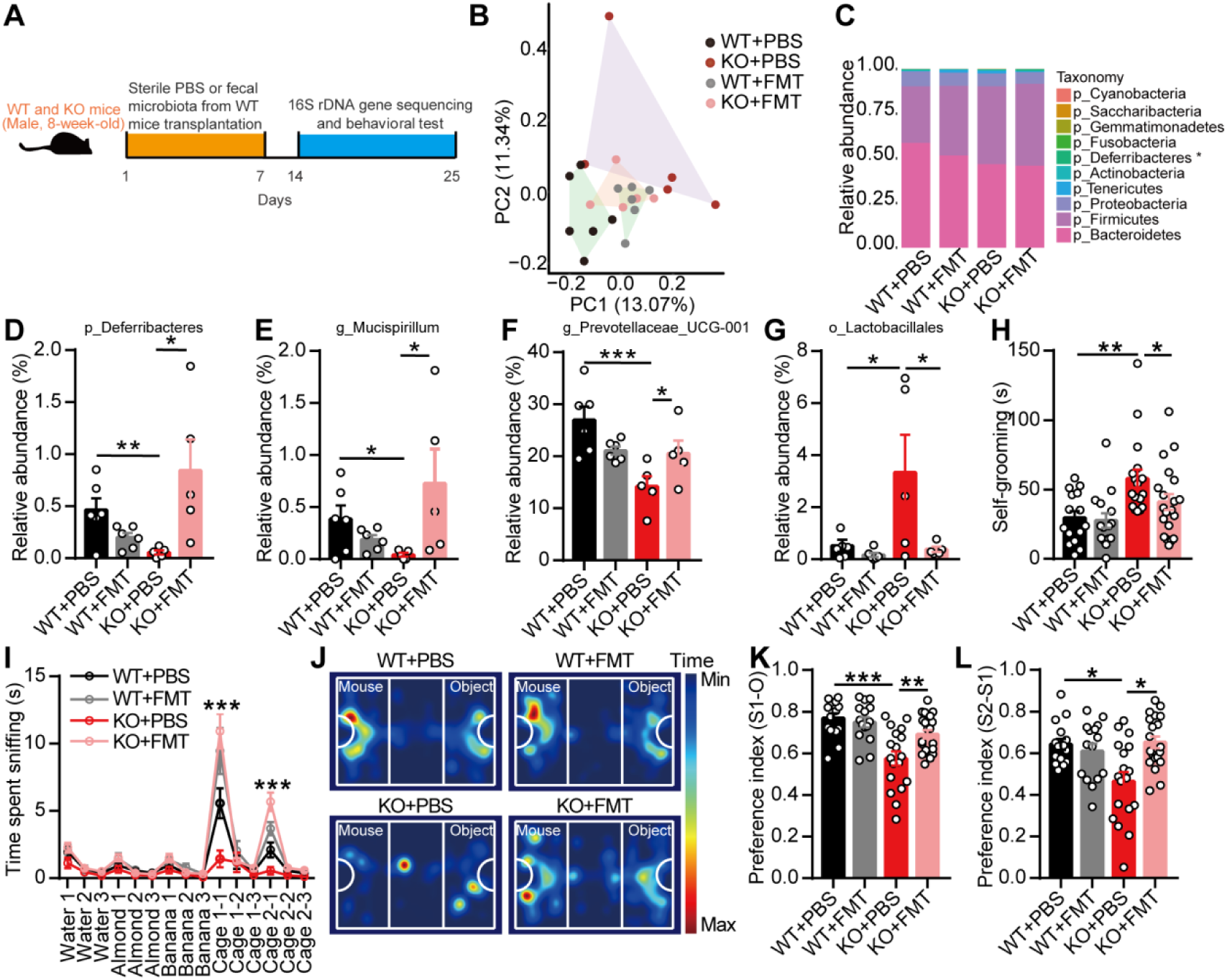
Transplantation of fecal microbiota from wild-type mice ameliorated autism-like behavior in adult mice with deletion of EphB6. (A) Schematic of the fecal microbiota transplantation. 8-week-old male WT and KO mice were gavaged orally with fecal microbiota of 8-week-old male WT mice (8 healthy mice from at least 3 cages) or sterile PBS for a week. After a week, fecal microbiota of the treated WT and KO mice were sequenced (B-G) and behavioral tests were conducted with an interval of at least 2 days (H-L). (B-G) 16S rDNA gene sequencing of fecal microbiota of mice. Principle coordinates analysis of Bray-Curtis distance (B). The relative abundance of top 10 bacteria in phylum level (C). In phylum level, the relative abundance of *Deferribacteres* (D). In species level, the relative abundance of *Mucispirillum* (E), *Prevotellaceae_UCG-001* (F) and *Lactobacillales* (G). n = 6, 6, 5, 5 mice respectively (H-L) Self-grooming test (H), olfactory habituation/dishabituation test (I), and three-chambered social approach task (J-L) were conducted. n = 15, 15, 18, 20 mice respectively. Data shown are mean ± SEM. One-way ANOVA (D-H, K-L), two-way repeated measures ANOVA (I), anosim analysis (B). *, p < 0.05; **, p < 0.01; ***, p < 0.001. WT, EphB6^+/+^ mice; KO, EphB6^−/−^ mice; FMT, fecal microbiota transplantation; PBS, phosphate buffered saline. Statistical values are presented in Supplementary Table 2.

Then functionally, we found, after being gavaged with fecal microbiota from WT mice, KO mice showed decreased self-grooming (Fig. 4H) and increased social behavior (Fig. 4I-L). These results indicated that gut microbial dysbiosis was responsible for autism-like behavior in mice with deletion of EphB6.

### Gut microbiota-mediated vitamin B6 metabolism regulated social behavior in EphB6-deficient mice

Considering the abnormal behaviors were probably because of the problem of brain, so we tried to figure out how gut microbiota affected brain and subsequently caused autism-like behavior in EphB6-deficient mice.

First, we tried to find the key region of brain affected by dysregulated gut microbiota in mice with deletion of EphB6, which was responsible for autism-like behavior. Studies on ASD patients or mouse models show that hippocampus, cerebellum and mPFC have been implicated in ASD (W. Cao et al., 2018; Rojas et al., 2006). After being processed with three-chambered social approach task, we found the protein expression of c-Fos was significantly increased in mPFC of KO mice compared with WT mice (Supplementary Fig. 3A-C). ASD has been generally considered to be caused by an increased ratio of synaptic excitation and inhibition and ASD children exhibit elevations in resting state neuronal activity (Gkogkas et al., 2013). So, whether mPFC was modulated by gut microbiota in KO mice needed to be further investigated. Because mPFC tissue was too small for some experiments and we used PFC tissue of mice in our next study.

The first question we asked was whether the bacteria could modulate mPFC directly. Unfortunately, we did not detect bacterial DNA or any bacterial colonies in PFC tissues of WT or KO mice (Supplementary Fig. 3D-E). Unexpectedly, we found metabolites of gut microbiota from KO mice also induced social deficits in C57BL/6J mice (Supplementary Fig. 3F-K). Were there some substances had been affected by gut microbial dysbiosis that caused social deficits in KO mice?

To found the metabolites that had been significantly changed, we detected metabolites in target tissue, that is PFC of KO mice, using non-targeted metabolomics strategies. Surprisingly, the metabolites in PFC were significantly different between the two groups of mice using orthogonal partial least squares discriminant analysis (Fig. 5A). KEGG pathway analysis showed 4 pathways that were significantly enriched in the differentially changed metabolites, including vitamin B6 metabolism pathway because of decreased relative abundance of pyridoxamine (PM) and pyridoxal 5’-phosphate (PLP) in PFC of KO mice (Fig. 5B-D).

**Fig. 5.**
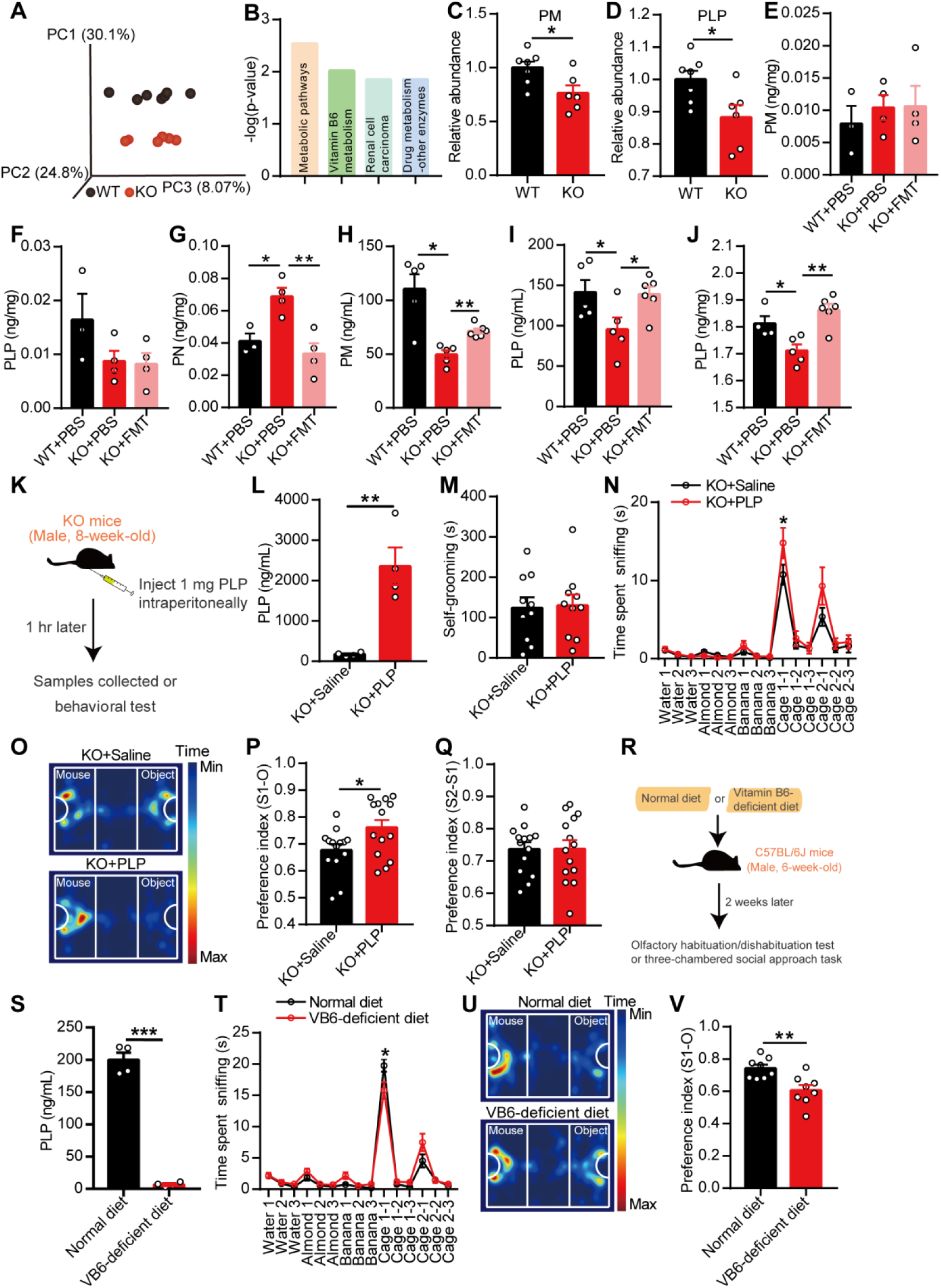
Gut microbiota-mediated vitamin B6 metabolism regulated social behavior in EphB6-deficient mice. (A-D) In non-targeted metabolomics analysis, the metabolites in PFC of 8-week-old male WT and KO mice were differently clustered by orthogonal partial least squares discriminant analysis (A). The enriched KEGG pathways associated with differential metabolites (B), the relative abundance of pyridoxamine (PM, C) and pyridoxal 5’-phosphate (PLP, D) were showed. n = 7, 6 mice respectively. (E-G) The level of PM (E), PLP (F) and pyridoxine (PN, G) in feces of 8-week-old KO mice gavaged with fecal microbiota from 8-week-old WT mice or PBS. n = 3, 4, 4 mice respectively. (H-J) The level of PM and PLP in plasma (H-I, n = 5, 5, 6 mice respectively) and level of PLP in PFC (J, n = 4, 5, 6 mice respectively) of 8-week-old KO mice gavaged with fecal microbiota from 8-week-old WT mice were increased compared with KO mice gavaged with PBS. (K-Q) Schematic of the injection of PLP (K). 8-week-old male KO mice were injected with 1 mg PLP or saline intraperitoneally. After 1 hr, mice were either sacrificed to detect PLP in plasma (L, n = 4 mice for each group) or subjected to self-grooming test (M, n = 10 mice for each group), olfactory habituation/dishabituation test (N, n = 8, 9 mice respectively) and three-chambered social approach task (O-Q, n = 14 mice for each group). Different mice were used for different behavioral tests. (R-V) Schematic of deficiency of vitamin B6 in 6-week-old SPF male C57BL/6J mice (R). Normal diet contained 12 mg vitamin B6 and vitamin B6-deficient diet were provided for 6-week-old C57BL/6J mice for 2 weeks. The level of PLP in plasma of C57BL/6J was detected (S, n = 4 mice for each group) and olfactory habituation/dishabituation test (T, n = 6, 8 mice respectively) or three-chambered social approach task (U-V, n = 8, 8 mice respectively) were conducted. Data shown are mean ± SEM. Two-tailed unpaired student’s *t* test (L-M, P-Q, S, V), R (B-D), one-way ANOVA (E-J), two-way repeated measures ANOVA (N, T). *, p < 0.05; **, p < 0.01; ***, p < 0.001. WT, EphB6^+/+^ mice; KO, EphB6^−/−^ mice; FMT, fecal microbiota transplantation; PBS, phosphate buffered saline, PFC, prefrontal cortex; PM, pyridoxamine; PLP, pyridoxal 5’-phosphate; PN, pyridoxine; VB6, vitamin B6. Statistical values are presented in Supplementary Table 2.

Vitamin B6 in body is mainly from diet and gut bacteria’s synthesis and then is absorbed in intestine. Then we detected the level of vitamin B6 in feces, blood and PFC of mice. The increased level of pyridoxine (PN) in feces, decreased level of PM and PLP in plasma, and decreased level of PLP in PFC were found in EphB6-deficient mice (Fig. 5E-J). A week after being gavaged with fecal microbiota from WT mice, KO mice had decreased level of PN in feces, increased level of PM and PLP in plasma and increased level of PLP in PFC compared with KO mice gavaged with sterile PBS (Fig. 5E-J). These results indicated that gut microbiota regulated the level of vitamin B6 in feces, blood and PFC of mice.

Vitamin B6 has been confirmed to be effective in a lot of ASD patients. Then we wondered if vitamin B6 supplementation could ameliorate the autism-like behavior of KO mice. However, the intragastric supplementation of vitamin B6 did not ameliorate social deficits of KO mice (Supplementary Fig. 4A-B). While one hour after being injected with 1 mg PLP intraperitoneally, KO mice had increased level of PLP in plasma (Fig. 5L), and increased social behavior (Fig. 5N-P) compared with control mice. While self-grooming (Fig. 5M) and social novelty (Fig. 5Q) were not changed in KO mice after the injection of PLP. However, the injection of 1 mg or 2 mg PLP intraperitoneally had no effect on social behavior of C57BL/6J mice (Supplementary Fig. 4C-E). Moreover, after being fed without vitamin B6 for two weeks, C57BL/6J mice had decreased level of PLP in plasma and decreased social behavior (Fig. 5R-V). Conclusively, our results hinted that there was a relation between gut microbiota-mediated defects of vitamin B6 metabolism and social deficits in EphB6-deficient mice.

### Gut microbiota-mediated vitamin B6 metabolism regulated dopamine in PFC in EphB6-deficient mice

Then we tried to clarify how the decreased vitamin B6 induced social deficits in mice. Vitamin B6, as a co-factor, has been implicated in more than 140 biochemical reactions in cells, including biosynthesis and catabolism of amino acid and neurotransmitters (Parra, Stahl, & Hellmann, 2018). As the most important active substances in brain, we first detected the neurotransmitters in PFC of mice by high performance liquid chromatography (HPLC) and found similar levels of glutamate, GABA, glycine, aspartic acid, serine and glutamine, among WT and KO mice gavaged with sterile PBS or fecal microbiota from WT mice (Fig. 6A). Interestingly, we found a decreased dopamine and an increased 5-HT in PFC of KO mice compared with WT mice (Fig. 6B). When treated with fecal microbiota from WT mice, KO mice had an increase in level of dopamine, but similar level of 5-HT, in PFC compared with KO mice gavaged with sterile PBS. While the level of noradrenaline, epinephrine and DOPAC were similar among the three groups of mice. More excitingly, after being gavaged with fecal microbiota from KO mice, the level of dopamine in PFC of SPF C57BL/6J mice had a decrease compared with C57BL/6J mice gavaged with fecal microbiota from WT mice (Supplementary Fig. 5A-C). Also, we found the injection of PLP intraperitoneally increased the level of dopamine in PFC of KO mice (Fig. 6C) and deficiency of vitamin B6 decreased the level of dopamine in PFC of SPF C57BL/6J mice (Fig. 6D). Briefly, these results indicated that gut microbiota-mediated vitamin B6 metabolism could affect the level of dopamine in PFC of mice.

**Fig. 6.**
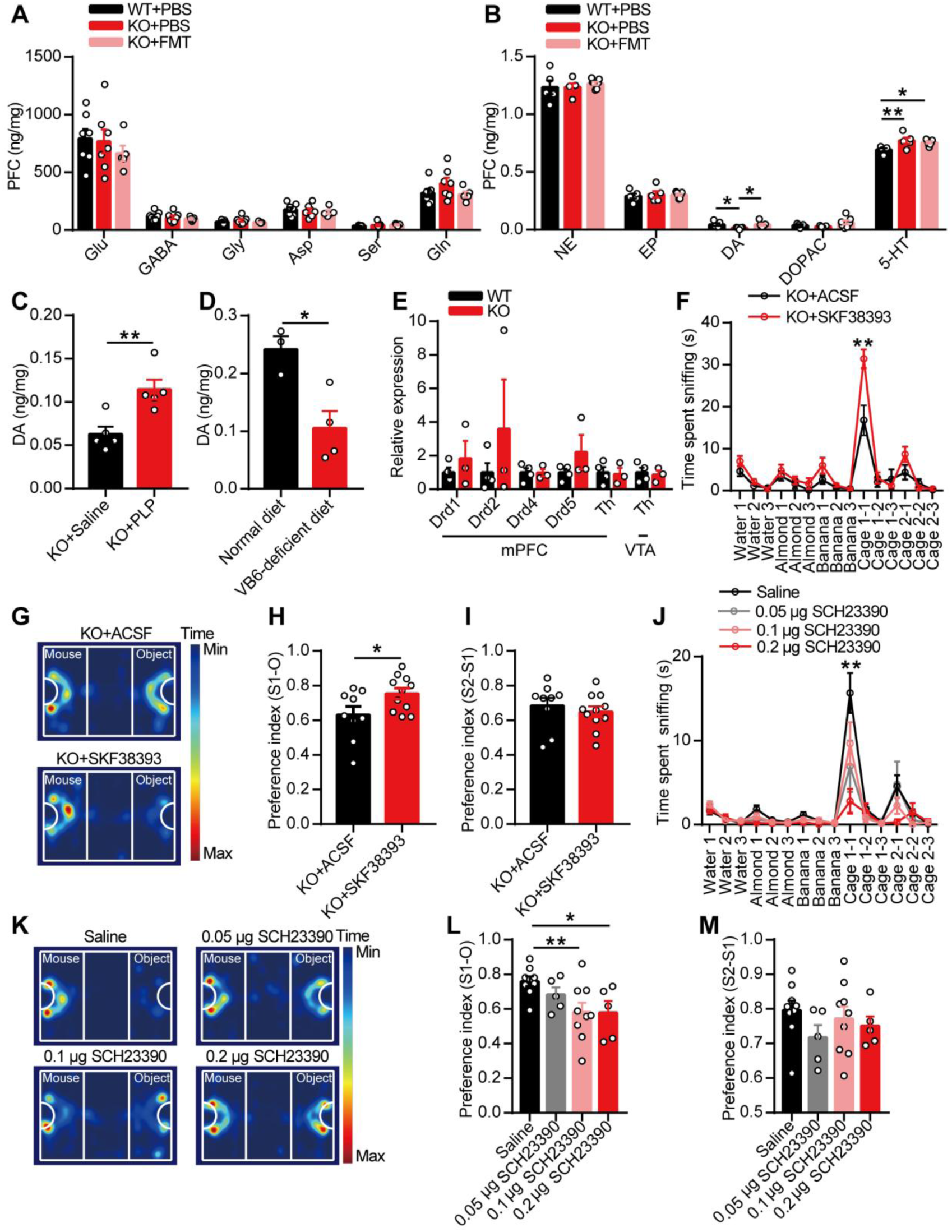
Dopamine was modulated by gut microbiota–mediated vitamin B6 and then regulated social behavior of EphB6-deficient mice. (A-B) The level of amino acid neurotransmitters (A, n = 7, 7, 5 mice respectively) and monoamine neurotransmitters (B, n = 5, 5, 7 mice respectively) in PFC of 8-week-old male WT and KO mice gavaged with sterile PBS or fecal microbiota from 8-week-old male WT mice were detected using HPLC. (C) 1 hr after the injection of 1 mg PLP intraperitoneally, the level of DA in PFC of 8-week-old male KO mice was increased compared with KO mice injected with saline. n = 5 mice for each group. (D) The level of DA in PFC of SPF male C57BL/6J mice fed without vitamin B6 was decreased compared with C57BL/6J mice fed with normal diet. n = 3, 4 mice respectively. (E) The mRNA expression of dopamine receptors and tyrosine hydroxylase in mPFC or VTA of 8-week-old WT and KO mice were detected by qRT-PCR. n = 4, 3 mice respectively. (F-I) Olfactory habituation/dishabituation test (F) and three-chambered social approach task (G-I) were conducted in 8-week-old male KO mice injected with D1R agonist (SKF38393, 0.0625 μg/0.3 μL) in mPFC with an interval of a week. n = 9, 11 mice respectively. (J-M) Olfactory habituation/dishabituation test (J, n = 8, 8, 9, 8 mice respectively) and three-chambered social approach task (K-M, n = 9, 5, 9, 5 mice respectively) were conducted in 8-week-old SPF male C57BL/6J mice injected with D1R antagonist (SCH23390) in mPFC with an interval of a week. Data shown are mean ± SEM. Two-tailed unpaired student’s *t* test (C-E, H-I), one-way ANOVA (A-B, L-M), two-way repeated measures ANOVA (F, J). *, p < 0.05; **, p < 0.01. WT, EphB6^+/+^ mice; KO, EphB6^−/−^ mice; FMT, fecal microbiota transplantation; PBS, phosphate buffered saline; PFC, prefrontal cortex; PLP, pyridoxal 5’-phosphate; VB6, vitamin B6; Glu, glutamic acid; GABA, gamma-aminobutyric acid; Gly, glycine; Asp, aspartic acid; Ser, serine; Gln, glutamine; NE, norepinephrine; EP, epinephrine; DA, dopamine; 5-HT, 5-hydroxytryptamine; DOPAC, dihydroxy-phenyl aceticacid; mPFC, middle prefrontal cortex; VTA, ventral tegmental area; Th, tyrosine hydroxylase; ACSF, artificial cerebrospinal fluid. Statistical values are presented in Supplementary Table 2.

To answer whether decreased dopamine contributed to autism-like behavior in EphB6-deficient mice and considering the fast metabolism of dopamine in brain, we injected the agonists of dopamine receptors into mPFC of mice. While deletion of EphB6 had no effect on mRNA expressions of dopamine receptors or tyrosine hydroxylase (Th) in mPFC or ventral tegmental area (VTA) (Fig. 6E). As dopamine D1 receptor (D1R) had the highest expression in mPFC and then followed by dopamine D2 receptor (D2R) (data not shown), we injected the D1R agonist (SKF38393) or D2R agonist (quinpirole) into mPFC of mice. We found KO mice had increased social behavior (Fig. 6F-I) after being injected with SKF38393 compared with KO mice injected with artificial cerebrospinal fluid (ACSF). While there were not any differences in C57BL/6J mice injected with ACSF or SKF38393 (Supplementary Fig. 5D-F). Differently, quinpirole did not increase social behavior in KO mice (Supplementary Fig. 5G-I). What’s more, D1R antagonist induced a decreased social behavior in C57BL/6J mice (Fig. 6J-M). In short, these results proposed dysregulated gut microbiota and vitamin B6 metabolism led to autism-like behavior by D1Rs-mediated pathway in EphB6-deficient mice.

### Gut microbiota regulated E/I balance in mPFC of EphB6-deficient mice

Imbalance between excitation and inhibition in synaptic transmission and neural circuits have been implicated in ASD (Cornew, Roberts, Blaskey, & Edgar, 2012; Rubenstein & Merzenich, 2003; Uhlhaas & Singer, 2012). E/I imbalance is frequently observed in animal models of ASD, and its correction can normalize key autistic phenotypes in these animals (Yizhar et al., 2011). Moreover, D1Rs are generally considered to modulate GABAergic inhibition in PFC (Seamans, Gorelova, Durstewitz, & Yang, 2001).

To further investigate the cellular mechanism underlying gut microbiota-mediated autism-like behavior in EphB6-deficient mice, we recorded spontaneous excitatory postsynaptic currents (sEPSCs) and spontaneous inhibitory postsynaptic currents (sIPSCs) of mPFC pyramidal neurons, in WT and KO mice treated with sterile PBS or fecal microbiota from WT mice. The amplitude and frequency of sEPSCs were similar among the groups (Fig. 7A-E). The amplitude of sIPSCs was also similar among the groups, while the frequency of sIPSCs was decreased in KO mice and was rescued by transplantation of fecal microbiota from WT mice (Fig. 7F-J). Additionally, we found a decreased frequency of sIPSCs in pyramidal neurons of mPFC in C57BL/6J mice gavaged with fecal microbiota of KO mice (Supplementary Fig. 6A-H). Collectively, these results indicated that gut microbiota modulated E/I balance, which was possibly regulated by dopamine, in pyramidal neurons of mPFC in EphB6-deficient mice.

**Fig. 7.**
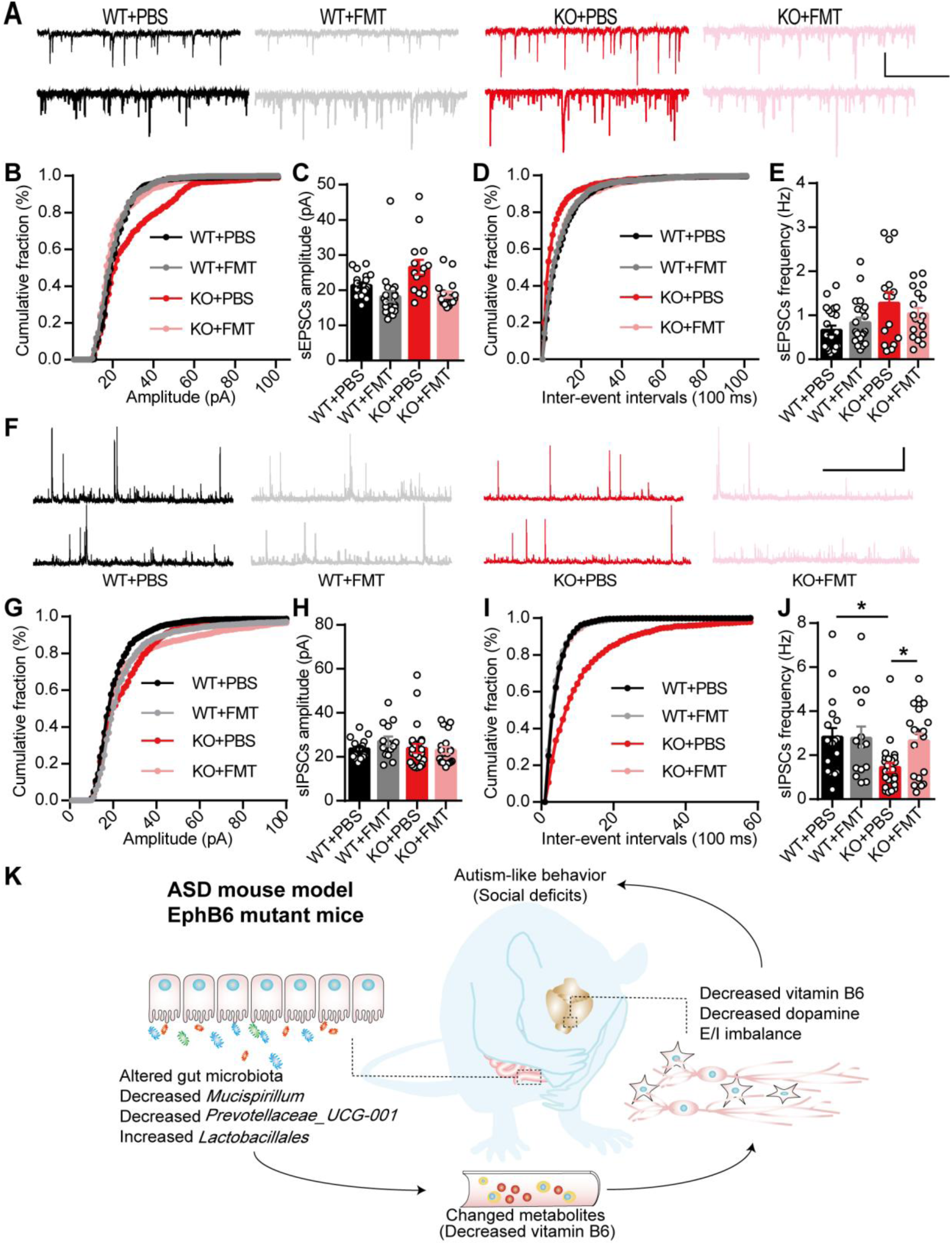
Fecal microbiota transplantation from wild-type mice ameliorated E/I imbalance in mPFC of EphB6-deficient mice. (A-E) Representative sEPSCs traces from pyramidal neurons in mPFC of 8-week-old male WT and KO mice gavaged with sterile PBS or fecal microbiota from 8-week-old WT mice (A), scale bars: 3 s, 20 pA. Cumulative distribution of sEPSCs amplitudes (B), average amplitude of sEPSCs (C). Cumulative distribution of sEPSCs frequencies (D), average frequency of sEPSCs (E). n = 22, 21, 15, 18 cells from at least 4 mice respectively. (F-J) Representative sIPSCs traces from pyramidal neurons in mPFC of 8-week-old male WT and KO mice gavaged with sterile PBS or fecal microbiota from 8-week-old WT mice (F), scale bars: 3 s, 20 pA. Cumulative distribution of sIPSCs amplitudes (G), average amplitude of sIPSCs (H). Cumulative distribution of sIPSCs frequencies (I), average frequency of sIPSCs (J). n = 18, 14, 25, 20 cells from at least 4 mice respectively. (K) The working model of the modulated social deficits in EphB6-deficient mice by gut microbiota. Deletion of EphB6 induced gut microbial dysbiosis and decreased vitamin B6 in plasma and PFC, which led to the decreased dopamine, E/I imbalance and social deficits in mice. Data shown are mean ± SEM. One-way ANOVA (C, E, H, J). *, p < 0.05. WT, EphB6^+/+^ mice; KO, EphB6^−/−^ mice; FMT, fecal microbiota transplantation; PBS, phosphate buffered saline; mPFC, middle prefrontal cortex; sEPSCs, spontaneous excitatory postsynaptic currents; sIPSCs, spontaneous inhibitory postsynaptic currents. Statistical values are presented in Supplementary Table 2.

## Discussion

Genomic studies for ASD patients based on two-generation sequencing strategies discover many candidate genes for ASD. Besides, increasing evidences, especially the clinical studies in ASD patients, suggest a functional link between gut microbiota and development of ASD. However, the ASD-associated genes and the mechanisms linking gut microbiota and brain dysfunctions (gut-brain axis) in ASD are still not well-established.

First in our study, we uncover EphB6 as an ASD-associated gene functionally. EphB6 has been suggested as a candidate gene for ASD for a long time (Agarwala, Shyamala, Padakannaya, & Ramachandra, 2019; Chen, 2009; O’Roak et al., 2011). Here, using our transgenic mouse models, we found deletion of EphB6 induced autism-like behavior in mice that mimicked the core symptoms of ASD patients fairly well. Using whole-genome sequencing, researchers have found more than 1000 genes that are associated with ASD, including EphB6 (Agarwala et al., 2019; Chen, 2009; O’Roak et al., 2011), EphA1 (Chen, 2009), EphB2 (Sanders et al., 2012). Our study uncovers the functional role of EphB6 in ASD and suggests EphB6-deficient mice can be used as a new mouse model of ASD.

Secondly, we find gut microbial dysbiosis is required for autism-like behavior in EphB6-deficient mice. Most ASD patients have serious GI problems (Buie et al., 2010; Coury et al., 2012; Parracho et al., 2005) and changed composition of gut microbiota (Gondalia et al., 2012; Strati et al., 2017). Moreover, microbiota transfer therapy can improve GI and autistic symptoms in ASD patients (D.-W. Kang et al., 2017; D. W. Kang et al., 2019). Our study suggests the probable role of gut microbiota on treating core symptoms of ASD patients, even in adult ones. Eph families have the important role in regulating epithelial homeostasis by the interaction with epithelial cell adhesion and junction proteins (Miao & Wang, 2009). Cldn4 can interact with EphA2 and ephrin-B1 to affect the tight junction integration (Tanaka, Kamata, & Sakai, 2005a, 2005b). AF-6 can be recruited to cell–cell contacts in MDCK and 293T cells by interacting with Eph receptors, including EphB6 (Buchert et al., 1999; Hock et al., 1998). The ablation of EphB6 may induce the dysregulated interaction between Eph families and junction proteins that leads to increased intestinal mucosal permeability and then gut microbial dysbiosis in mice.

Thirdly, we find dysfunction of vitamin B6 metabolism is key for gut microbiota-mediated autism-like behavior in EphB6-deficient mice. We found the decreased level of vitamin B6 in plasma and PFC of EphB6-deficient mice was rescued by transplantation of fecal microbiota from wild-type mice. Moreover, injection of vitamin B6 intraperitoneally rescued social deficits of EphB6-deficient mice. More interestingly, PLP has been found to have an unbelievable low level in ASD children compared with controls (Adams & Holloway, 2004). Date to 1960s, a lot of clinical studies have used vitamin B6 to treat ASD children, and most studies have reported the improved autistic symptoms in ASD children (Mousainbosc et al., 2006; Rossignol, 2009; Sato, 2018). While there are also reports of the non-effect of vitamin B6 on ASD patients (Murza, Pavelko, Malani, & Nye, 2010). Considering the complicated causes of ASD, we think vitamin B6 is effective for a part of ASD patients, such as ASD patients with EphB6 mutation. Vitamin B6 cannot be synthesized by the body itself and the source of vitamin B6 in body is mainly from diet and bacteria’s synthesis via intestinal absorption. So, normal intestinal functions are important for the homeostasis of vitamin B6 in body. The intestinal absorption of vitamin B6 is pH dependent with higher uptake at acidic compared with alkaline pHs (Said, Ortiz, & Ma, 2003). The more alkaline environment in gut of EphB6-deficient mice (Supplementary Fig. 1I) may cause decreased absorption of vitamin B6. How the changed bacteria affect the gut pH and vitamin B6 level in blood need more exploration. Overall, our study finds a new modulated role of gut microbiota on vitamin B6 and proves the ameliorative role of gut microbiota-mediated vitamin B6 on social deficits in EphB6-deficient mice.

Finally, we functionally establish the mechanisms linking gut microbiota and brain dysfunctions (gut-brain axis) in EphB6-deficient mice. Gut-brain axis has been generally considered to be involved in psychiatric diseases. However, there are few studies on how brain is specially regulated by gut microbiota. In our study, we found dopamine in PFC of EphB6-deficient mice was regulated by gut microbiota-mediated vitamin B6. In ASD patients, there are lower medial prefrontal dopaminergic activity (Ernst, Zametkin, Matochik, Pascualvaca, & Cohen, 1997). After being given vitamin B6, autistic children have a reduced urinary homovanillic acid, which suggests an improved dopamine metabolism (Lelord et al., 1978). These studies suggest the regulated role of vitamin B6 in dopaminergic metabolites in ASD patients. Previously, Sgritta reported the modulated VTA plasticity by *Lactobacillus reuteri* in ASD mouse models (Sgritta et al., 2019). Here, our study showed a new regulatory role of gut microbiota on dopamine in PFC by modulating vitamin B6 in EphB6-deficient mice. What’s more, we found the ameliorative role of D1R agonists in social behavior and the modulated E/I balance in mPFC by gut microbiota in EphB6-deficient mice. Activating D1Rs in PFC can increase frequency of sIPSCs of pyramidal neurons, while D2R agonist does not have the same effect (Seamans et al., 2001). The modulation of social behavior by D1Rs was probably because of its modulation of GABAergic inhibition in EphB6-deficient mice. Collectively, our study indicates decreased dopamine is induced by dysregulated gut microbiota-mediated defect of vitamin B6 and then contributes to E/I imbalance and social deficits in EphB6-deficient mice.

In summary, our study uncovers a key role of EphB6 in autism-like behavior. Mechanistically, gut microbiota-mediated defect of vitamin B6 metabolism regulates autistic-like social behavior by inducing E/I imbalance in mPFC in EphB6-deficient mice. Our study suggests a new ASD mouse model and provides a new insight into gut-brain axis.

## Materials and methods

### Mice

All the mice used in our experiments were male. SPF C57BL/6J mice (3-8 weeks old) were obtained from Animal Experiment Center of Southern Medical University in Guangzhou of China. EphB6-deficient mice were obtained using the embryonic stem (ES) cells that were inserted with EphB6^tm1e(KOMP)Wtsi^ vector (IKMC project number: 49365). The injection of ES cells and obtaining of chimeric mice were operated by subsidiary of Cyagen Biosciences Inc. in Guangzhou of China. The chimeric mice were crossed with SPF C57BL/6J mice and its offspring (EphB6^+/−^ mice) were kept being crossed with SPF C57BL/6J mice until the fifth generation was born. Then EphB6^+/−^ mice were crossed with each other to obtain EphB6^+/+^ mice and EphB6^−/−^ mice. All the mice were raised at a controlled appropriate SPF condition with the temperature at 24 ± 1°C and humidity at 50% to 70% separately and with 12 hr light-dark cycles by turning lights on from 8:00 a.m. to 8:00 p.m.. The animals were housed in groups of 4-5 in plastic cages (Exhaust Ventilated Closed-System Cage Rack). Standard sterile diet and drinking water for raising mice were used to feed the mice. The EphB6 ablation mice were genotyped at 2-week-old using the primers 5’-CTCTGCCAAGGTGAGACACTTTTCC-3’ and 5’-AGCCAGTCTCTACCTCCTGTTTTGG-3’ for the wild-type band, 5’-CTCTGCCAAGGTGAGACACTTTTCC-3’ and 5’-CGTGGTATCGTTATGCGCCT-3’ for the mutant band and weaned at 3-week-old. All procedures treated with mice were in compliance with the Regulations for the Administration of Affairs Concerning Experimental Animals in China.

### Fecal microbiota transplantation

Fresh feces of healthy male EphB6^+/+^ and EphB6^−/−^ mice (8 mice for each group from at least 3 cages) were collected from the disinfected anus into new sterile tubes every day before the experiment to promise microbial vitality (Olson et al., 2018). Then the fresh feces were weighted, mixed with sterile PBS at a dilution ratio of 1 mg/10 μL or 1 mg/20 μL and centrifuged at 900 × g for 3 min. The supernatant was collected and gavaged orally to each mouse (10 mL/kg) for 5 or 7 consecutive days. All the mice were handled aseptically.

The dilution ratio of 1 mg/20 μL was used to treat 3-week-old SPF C57BL/6J mice and the dilution ratio of 1 mg/10 μL was used to treat 6-week-old SPF C57BL/6J mice and 8-week-old EphB6^+/+^ and EphB6^−/−^ mice.

For fecal microbiota or metabolite transplantation, after the fresh feces were weighted, mixed with sterile PBS at a dilution ratio of 1 mg/10 μL and centrifuged at 4000 × g for 10 min, the supernatant and precipitate were both collected. After being filtered by the filter with a pore size of 0.22 μm (Cat# SLGP033RS, Millipore, Darmstadt, Germany), the supernatant was orally gavaged to each mouse (10 mL/kg) for 7 consecutive days. After being resuspended in sterile PBS, centrifuged at 900 × g for 3 min, washed by sterile PBS twice, the precipitate was orally gavaged to each mouse (10 mL/kg) for 7 consecutive days.

### Antibiotics treatment

Vancomycin (50 mg/kg, CAS: 123409-00-7, MP bio, California, USA), neomycin (100 mg/kg, CAS: 1405-10-3, MP bio) and metronidazole (100 mg/kg, CAS: 443-48-1, MCE, New Jersey, USA) were mixed using sterile drinking water (Olson et al., 2018). Then the mixture was orally gavaged to 3-week-old or 6-week-old SPF C57BL/6J mice twice a day for 5 or 7 consecutive days and the amount of infusion was based on the weight of mice. The mixture was prepared every day and used freshly. During the treatment, ampicillin (1 mg/mL, CAS: 69-52-3, MP bio) was added into the drinking water of mice and changed with fresh solution every 3 days. For 3-week-old SPF C57BL/6J mice, the antibiotics treatment lasted for 5 days (Gong et al., 2018). All the mice were handled aseptically.

### Western blot analysis

After abdominally anesthetized with phenobarbital sodium (60 mg/kg), the brain of mouse was quickly removed, put into an ice-cold mouse brain mold (Cat# 68713, RWD, Shenzhen, China) and sliced. Then posterior mPFC, hippocampus and cerebellum of mice were cut out. The total proteins of tissues were extracted using the lysis buffer (Cat# P0013B, Beyotime, Shanghai, China) and boiled in protein loading buffer. Equal amounts of the denatured protein samples were electrophoresed in 6-10% polyacrylamide gel containing 0.1% SDS and transferred to polyvinylidene fluoride (PVDF) membranes with pore size of 0.45 μm (Cat# IPVH00010, Millipore). Then the PVDF membranes were incubated with primary antibodies at 4°C for at least 12 hr. After that, the samples were incubated with secondary antibodies for about 2 hr at room temperature (Cat# BA1050 and Cat# BA1054, Boster, California, USA). The desired signals were visualized by Quantitative FluorChem SP Imaging System (Alpha Innotech, California, USA). Intensities of bands were quantified by ImageJ software and signal values of corresponding band of Gapdh were considered as internal controls. The following primary antibodies were used: EphB6 (1:500, Cat# ab54656, Abcam, Cambridge, UK), c-Fos (1:500, Cat# MABE329, Millipore) and Gapdh (1:5000, Cat# 60004-1-ig, Proteintech, Chicago, USA).

### Quantitative reverse transcription PCR (qRT-PCR)

After anesthetized with phenobarbital sodium (60 mg/kg), different tissues of mice were quickly removed and put into liquid nitrogen, including colon, colonic epithelium, spleen and lung. Then posterior mPFC and VTA were sectioned out using ice-cold mouse brain mold (Cat# 68713, RWD). qRT-PCR was performed accordingly (Ying Li et al., 2017) by using a 7500 real-time PCR system (ABI, California, USA) and SYBR Premix Ex Taq (Cat# RR420A, Takara, Osaka, Japan). Normalized to the mRNA expression level of Gapdh or Actb, the mRNA expressions of other genes were evaluated using the method of ΔΔCt. All primers used in qRT-PCR were listed in Supplementary Table 1.

### Hematoxylin-eosin staining

The staining of different tissues of mice with haematoxylin and eosin was performed accordingly (Ying Li et al., 2017). Briefly, the tissues were immersed into 4% formaldehyde immediately for 24 hr. Then tissues were embedded in paraffin, sectioned, and stained with haematoxylin and eosin. The stained sections were observed using an optical microscope (Olympus, Tokyo, Japan).

### Intestinal permeability assay

Mice were fasted for 4 hr before experiment, then FITC-dextran (50 mg/mL, Cat# 46944, Sigma Aldrich, Missouri, USA) was gavaged to mice (600 mg/kg) (Hsiao et al., 2013). 4 hr after the oral gavage, the blood of mouse was collected by cardiac puncture. Then the blood was placed at room temperature for 1 hr before being centrifuged at a speed of 3000 rpm for 10 min. Then the supernatant was transferred to a new tube and centrifuged at a speed of 12000 rpm for 10 min at 4°C. The supernatant, which was the serum, was diluted with equal volume of PBS and 100 μL diluted serum was added to a 96-cell microplate. The concentration of FITC in serum was determined by Varioskan LUX microplate reader (Thermo Fisher Scientific, Massachusetts, USA) with an excitation of 485 nm and an emission wavelength of 528 nm. The serial diluted FITC-dextran (0, 0.5, 1, 2, 4, 6, 8, 10 μg/μL) was used as standards. Serum of mice administered with PBS was used as negative controls.

### Vitamin B6-deficient mouse model

The formula of diet with normal vitamin B6 or without vitamin B6 was based on previous study (Qian, Shen, Zhang, & Jing, 2017). Then 6-week-old SPF male C57BL/6J mice were fed with the diet with or without vitamin B6 for two weeks.

### Behavioral studies

Mice used for experiments were male and naive. Mice were handled for 3 days before the experiments and habituated in the experiment room for at least 30 min before each test (R. M. Deacon, 2006). Mice were performed with different behavioral tests with a sequence or different mice were used for different behavioral tests which were mentioned in the figure legends. The sequence of different behavioral tests was self-grooming test, olfactory habituation/dishabituation test, three-chambered social approach task, marble burying test, open field test, social partition test, elevated plus maze and morris-water-maze test. Different behavioral tests were done with an interval of at least 2 days.

Self-grooming test was performed as previously described (McFarlane et al., 2008). Generally, mouse was first placed in an empty crystal cage to habituate the cage for 10 min, then the time each mouse spent on self-grooming was recorded during next 10 min by a double-blind experienced experimenter. Self-grooming included face-wiping, scratching/rubbing of head and ears, and full-body grooming. Between each trial, the apparatus was cleaned by 30% ethyl alcohol in water.

Marble burying test was performed as previously described (Robert MJ Deacon, 2006). Mouse was placed into an animal cage filled with fresh wood chip bedding with the depth of 5 cm. Regular pattern of glass marbles (5 rows of 4 marbles), which were placed 4 cm apart from each other, were regularly placed under the bedding and mouse was allowed to explore for 30 min. The number of buried (more than 50 percent of their depth in bedding) marbles was counted.

Social partition test was performed as previously described (Lugo, Swann, & Anderson, 2014). Mouse was individually housed in one side of the cage which was divided by a clear perforated partition with 0.6 cm-diameter holes, and the other side of cage was housed with a sex- and age-matched C57BL/6J mouse for 24 hr before experiment. At the first trial, the total time that the experimental mouse spent on sniffing partition with the familiar mouse on the other side during 5 min was recorded. Then the familiar mouse was replaced with a sex- and age-matched unfamiliar C57BL/6J mouse, the total time that the experimental mouse spent on sniffing partition with the unfamiliar mouse during 5 min was recorded. In the last trial, unfamiliar mouse was instead replaced by familiar mouse and the time that the experimental mouse spent on sniffing partition as first trial was recorded. The time spent on sniffing partition in the three trials was recorded by a double-blind experienced experimenter.

Olfactory habituation/dishabituation test was performed as previously described (Silverman, Yang, Lord, & Crawley, 2010). Mouse was placed into a clean usual animal cage with thin bedding in a fresh room for 30 min before test. One swab saturated with water was given to mouse for 2 min, and then quickly replaced by another swab saturated with water for the following 2 min, then third swab saturated with water was given to mice for another 2 min quickly. Then other odors were given to mouse similarly. The sequence of given odors was water, almond extract, imitation banana flavor, odor of soiled bedding from mice and odor of soiled bedding from another cage of mice. Water, almond extract (dilution of 1:100) and imitation banana flavor (dilution of 1:100) were regarded as unsocial odors, while soiled bedding with the excrement of sex- and age-matched unfamiliar C57BL/6J mice were regarded as social odors. The time each mouse spent on sniffing the odorant swabs in every 2 min trial was recorded by a double-blind experienced experimenter.

Three-chambered social approach task was performed as previously described (Yang, Silverman, & Crawley, 2011). The apparatus was divided into three rectangular clear chambers (60 cm × 40 cm × 22 cm) by two walls on which had two removable doorways (8 cm × 5 cm) that allowed mouse to access each chamber freely. After habituated to the middle chamber for 5 min, the mice were allowed to explore the three chambers freely for 10 min. For the sociability test, an age- and sex-matched C57BL/6J mouse was placed in the wire cage in one chamber while the wire cage in the other chamber was empty. Then the dividers were raised and the experimental mouse was allowed to freely explore all three chambers for 10 min. For the social novelty test, another age- and sex-matched C57BL/6J mouse was placed in the empty wire cage described above. And the experimental mouse was originally placed in the center of the chamber and allowed to explore freely for 10 min after doorways were removed. Between each trial, the apparatus was cleaned by 70% ethyl alcohol in water. Time spent in each chamber and heat maps were calculated using EthoVison XT software (Noldus, Wageningen, Netherlands). And the time spent on sniffing the wire cages which represented the social approach behavior of mice was calculated by a double-blind experienced experimenter.

In open field test, mouse was placed in the center of an open field chamber (40 cm × 40 cm × 30 cm) (X. Cao et al., 2013). Exploratory behavior of mice was assessed by a session of 30 min and total distance was automatically recorded and analyzed by a VersaMax animal behavioral monitor system (Omnitech Electronics, Nova Scotia, Canada).

Elevated-plus-maze test was performed accordingly (X. Cao et al., 2013). Briefly, mouse was put into the center of elevated-plus-maze which was consisted of two opposing open arms (30 cm × 5 cm × 0.5 cm), two opposing enclosed arms (30 cm × 5 cm × 15 cm) and a central platform (5 cm × 5 cm) for 5 min. The time spent in different arms and entries into different arms were recorded by EthoVison XT software (Noldus).

Morris water maze test was performed as before (Vorhees & Williams, 2006). Generally, 4 trials were given to each mouse every day for 5 days. In each trial, the searching time for the mouse was no more than 1 min. A stay on the platform was 15 s. Intervals between each trial were no less than 1 min. On the sixth day, the probe test was performed by removing the platform and recording the swimming paths of mice in 1 min. The swimming paths of mice during the learning and test period were analyzed by EthoVison XT software (Noldus).

### 16S rDNA gene sequencing

Fecal samples of the experimental mice were collected and stored at −80°C before being performed. Using QIAamp Fast DNA Stool Mini kit (Cat# 51604, QIAGEN, Venlo, Netherlands), genomic DNA of samples were extracted. The purity and concentration of the extracted DNA were detected using agarose gel electrophoresis. Bacterial DNA was amplified with the primers targeting V3-V4 regions (5’-TACGGRAGGCAGCAG-3’, 5’-GGGTATCTAATCCT-3’). Then DNA was sequenced using MiSeq PE300 platform (Illumina, California, USA) by oe biotechnology company in shanghai. The raw data were treated and processed using QIIME software package (version 1.8.0). Then represent sequences of OTU were blasted in Silva database (version 123). The alpha diversity and beta diversity were analyzed using QIIME software package (version 1.8.0).

### Metabolomic analysis

For untargeted metabolite analysis, plasma and PFC of mice were prepared and deproteinized with methanol. Then the samples were analyzed using liquid chromatography-mass spectrometry by oe biotechnology company in shanghai. UPLC-Q-TOF/MS (ACQUITY UPLC I-Class, Waters, Massachusetts, USA) and ESI-QTOF/MS (Xevo G2-S Q-TOF, Waters) were used. The chromatographic column was the ACQUITY UPLC BEH C18 Column (1.7 µm, 2.1 mm X 100 mm, Waters). Mobile phase A was water contained with 0.1% formic acid and mobile phase B was acetonitrile contained with 0.1% formic acid. The gradient elution was 1%-5% mobile phase B in 0-1 min, 5%-30% mobile phase B in 1-2 min, 30%-60% mobile phase B in 2-3.5 min, 60%-90% mobile phase B in 3.5-7.5 min, 90%-100% mobile phase B in 7.5-9.5 min, 100% mobile phase B in 9.5-12.5 min, 100%-1% mobile phase B in 12.5-12.7 min, 1% mobile phase B in 12.7-16 min. The spectrum signal of samples was acquired by electrospray ionization using positive and negative ionization modes. The data were pretreated using progenesis QI (Waters) and then multivariate statistical analysis was performed using SIMCA software (version 14.0, Umetrics, Umeå, Sweden). The enriched pathway analysis of changed metabolites was performed using KEGG database (http://www.genome.jp/KEGG/pathway.html) and R (version 3.4.1).

For targeted metabolic analysis, PFC was pretreated with 0.4 M perchloric acid which contained 0.04% EDTA and 100 µL plasma was pretreated with 50 µL 5% trichloroacetic acid. 1 M NaOH was added to samples to quench acid.

For the analysis of amino acid neurotransmitters, high performance liquid chromatography (HPLC, Shimadzu, Kyoto, Japan) was used, with fluorescence detection system (Prominence RF-20A/20Axs, Shimadzu) and the C18 chromatographic column (Eclipse AAA, 4.6 × 150 mm, 5 μm, Agilent, California, USA). All the used reagents and liquid were chromatographically pure. Mobile phase A contained 20 mM sodium acetate solution (pH7.2), methyl alcohol and tetrahydrofuran which were at a volume ratio of 400:95:5. Mobile phase B contained 20 mM sodium acetate solution (pH7.2) and methyl alcohol which were at a volume ratio of 120:380. The gradient elution was 0%-63% mobile phase B in 0-10 min, 63% mobile phase B in 10-12 min, 63%-100% mobile phase B in 12-12.01 min, 100% mobile phase B in 12.01-17 min, 100%-0% mobile phase B in 17-18 min, 0% mobile phase B in 18-21 min. The temperature of column was set as 35°C. The flowing rate of mobile phase was 0.8 mL/min. The excitation wavelength was set as 340 nm, and the emission wavelength was set as 455 nm. The derivatization reagent contained o-phthalaldehyde (OPA, 5 mg, CAS: 643-79-8, Sigma-Aldrich), methyl alcohol (120 μL), β-mercaptoethanol (10 μL) and borate buffer (0.2 M, pH 9.2, 1mL) and was kept out of light. Data were recorded by INT7. The standards, including glutamic acid (CAS: 56-86-0, Sigma-Aldrich), gamma-aminobutyric acid (CAS: 56-12-2, Sigma-Aldrich), glycine (CAS: 56-40-6, Sigma-Aldrich), aspartic acid (CAS: 56-84-8, Sigma-Aldrich), serine (CAS: 56-45-1, Sigma-Aldrich) taurine (CAS: 107-35-7, Sigma-Aldrich) and glutamine (CAS: 56-85-9, Sigma-Aldrich), were prepared at the concentrations of 7.8125, 15.625, 31.25, 62.5, 125 and 250 ng/mL. The concentrations of amino acid neurotransmitters in different samples were acquired according to the concentrations of standards.

For the analysis of monoamine neurotransmitters, HPLC contained with electrochemical detection system (DECADE life, Antec Scientific, Zoeterwoude, Netherlands) was used, including the chromatographic column (Accucore C18, 150 × 2.1 mm, 2.6 μm, Thermo Scientific). All the used reagents and liquid were chromatographically pure. Mobile phase was prepared with deionized water and MeOH with a volume ratio of 9:1, containing NaH_2_PO_4_ (100 mM), sodium octane sulfonate (0.74 mM), Na_2_EDTA (0.027 mM) and KCl (2 mM). The temperature of column was set at 35°C. And the flowing rate of mobile phase was set at 0.2 mL/min. The standards, including epinephrine (CAS: 51-43-4, Sigma-Aldrich), noradrenaline (CAS: 108341-18-0, Sigma-Aldrich), dopamine (CAS: 62-31-7, Sigma-Aldrich), 3,4-dihydroxyphenylacetic acid (CAS: 102-32-9, Sigma-Aldrich), homovanillic acid (CAS: 306-08-1, Sigma-Aldrich), 5-Hydroxyindole-3-acetic acid (CAS: 54-16-0, Sigma-Aldrich) and 5-hydroxytryptamine (CAS: 153-98-0, Sigma-Aldrich), were prepared at the concentrations of 0.5, 1, 25, 125 and 250 ng/mL. The concentrations of monoamine neurotransmitters in different samples were acquired according to the concentrations of standards.

For the analysis of pyridoxal 5’-phosphate and pyridoxamine, TSQ Quantiva combined with Prelude SPLC System (Thermo Fisher Scientific) were used. All the used reagents and liquid were chromatographically pure. First, the separation of substances was performed using Prelude SPLC System with the C18 chromatographic column (Water Acquity UPLC HSS T3, 2.1 × 100 mm, 1.7 μm). Mobile phase A contained 0.2% formic acid. Mobile phase B was methyl alcohol. The gradient elution was 0%-50% mobile phase B in 0-2 min, 50%-95% mobile phase B in 2-3.5 min, 95% mobile phase B in 3.5-5.5 min, 95%-0% mobile phase B in 5.5-6.5 min. The temperature of column was set as 40°C. The flowing rate of mobile phase was 0.25 mL/min. Data were recorded using positive-ion electrospray ionization and the selected reaction monitoring mode. For pyridoxal 5’-phosphate, precursor ion was *m/z* 248.03, product ion was *m/z* 150.071 and collision energy was 16.067 V. For pyridoxamine, precursor ion was *m/z* 169.152, product ion was *m/z* 152.111 and collision energy was 12.124 V. Data were acquired and processed with TraceFinder software (version 3.3 sp1, Thermo Fisher Scientific). The standards, including pyridoxal 5’-phosphate (CAS: 41468-25-1, Sigma-Aldrich) and pyridoxamine (CAS: 524-36-7, Sigma-Aldrich), were prepared at the concentrations of 0.1, 0.5, 1, 6.25, 12.5, 25, 50 and 100 ng/mL. The concentrations of pyridoxal 5’-phosphate and pyridoxamine in different samples were acquired according to the concentrations of standards.

### Bacterial culturing

PFC of mice were brought out aseptically and homogenized in PBS using sterile magnetic beads. Then the homogenates were painted on the Luria-Bertani solid medium and cultured for 24 hr at 37°C.

### DNA extraction of bacteria

The PFC of mice were brought out aseptically and the genomic DNA of the tissue was extracted using PureLink genomic DNA kit (Cat# K1820-01, Invitrogen, California, USA). Then the DNA was amplified using bacterial universal primers (5’-AGAGTTTGATCATGGCTCAG-3’, 5’-CCGGGAACGTATTCACC-3’) (Ueno et al., 2015). The DNA of *Escherichia coli* was used as positive control.

### Stereotaxic surgery and drug microinjection

The stereotaxic surgery was performed to implant brain infusion cannula into mPFC of adult male mice based on the published protocols (X. Cao et al., 2013). After being anesthetized by phenobarbital sodium (60 mg/kg), the mouse was placed in a stereotaxic frame (RWD) and a hole with the diameter of 1 mm was drilled with a dental drill on the skull of mouse according to the adjusted coordinates of bilateral mPFC (AP: +1.84 mm, ML: ± 0.4 mm, DV: −2.2 mm). Then the brain infusion cannula (Cat# 62004, RWD) was carefully put into the drilled hole and fixed by glass ionomer cement. After the operation, the mice were resuscitated on an electric blanket and then put back into the original cage. After a recovery of 7 days, the catheter cap (Cat# 62104, RWD) was removed and the injection needle (Cat# 6220, RWD4) was inserted into the catheter after a disinfection with 75% alcohol. The injection needle was connected with a microsyringe through a polyethylene tube (Cat# 62320, RWD), and the drug was injected into the mPFC at a speed of 0.1 μL/min controlled by a microsyringe pump (Cat# R404, RWD). The total volume of the injected drug was 0.3 μL. After the injection of drug, the injection needle was kept being inserted into the catheter for 5 min to facilitate the complete diffusion and absorption of the drug. Behavioral test was performed 30 min after administration. The drugs used in this experiment were SKF38393 (CAS: 62717-42-4, MCE) and quinpirole (CAS: 524-36-7, Sigma-Aldrich).

### Slice preparation

Male mice were decapitated after anesthetized by phenobarbital sodium (60 mg/kg). Brains were removed quickly and then placed into the ice-cold modified ACSF containing (in mM): 26 NaHCO_3,_ 10 glucose, 10 MgSO_4_, 2 KCl, 1.3 NaH_2_PO_4_, 0.2 CaCl_2_ and 250 sucrose. Slices containing mPFC (300 µm) were prepared using a VT-1200S vibratome (Leica, Wetzlar, Germany) in ice-cold modified ACSF. And then slices were transferred into the storage chamber containing the regular ACSF (in mM) (126 NaCl, 26 NaHCO_3_, 10 glucose, 2 CaCl_2_, 3 M KCl, 1 MgSO_4_, and 1.25 NaH_2_PO_4_) at 31°C for 1 hr and then were removed to room temperature (25 ± 1°C) for 1 hr before being recorded. All solutions were saturated with 95% O_2_/5% CO_2_ (vol/vol) during the slice preparation (Y. Li et al., 2018).

### Electrophysiology

The neurons in mPFC were obtained using an infrared (IR)-differential interference contrast (DIC) microscope (ECLIPSE FN1, Nikon, Tokyo, Japan). To record sEPSCs, pipettes (input resistance: 3-7 MΩ) were filled with an intracellular solution containing (in mM) 105 K-gluconate, 30 KCl, 10 phosphocreatine, 10 HEPES, 4 ATP-Mg, 0.3 EGTA, and 0.3 GTP-Na (pH 7.3, 285 mOsm). When recording sEPSCs, the GABA_A_ receptors were blocked with 20 µM bicuculline methiodide (CAS:40709-69-1, TOCRIS, Minneapolis, USA). When recording sIPSCs, the holding potentials were 0 mV, pipettes (input resistance: 3-7 MΩ) were filled with an intracellular solution containing (in mM) 110 Cs_2_SO_4_, 0.5 CaCl_2_, 2 MgCl_2_, 5 EGTA, 5 HEPES, 5 TEA, 5 ATP-Mg (pH 7.35, 285 mOsm). Data were recorded by a multiClamp 700B (Molecular Devices, San Jose, USA), digitized at 10 kHz and filtered at 3 kHz. Data were collected when the series resistance fluctuated within 20% of the initial values and analyzed by pClamp 10.2 software (Molecular Devices) (Y. Li et al., 2018).

### Statistical analyses

All statistical analyses were performed with SPSS statistical software (version 20.0). Sample size was determined according to previously published studies (Buffington et al., 2016; Hsiao et al., 2013; Olson et al., 2018; Sgritta et al., 2019). No animals were excluded. The normality of all data was analyzed using Kolmogorov-Smirnov test. Levene’s test was used for the test of equal variances. For the data with normal distributions, two-tailed and unpaired Student’s *t*-test was performed to analyze two independent groups with equal variance. Two-tailed and unpaired Student’s *t*-test with Welch’s correction was used to analyze two independent groups with unequal variance. One-way ANOVA was performed to analyze multiple groups with only one variable and the differences between groups were performed with LSD pairwise comparison. When the variances were unequal, Dunnett’s T3 pairwise comparison was used to analyze the difference between groups. For the data without normal distributions, Mann-Whitney *U* test was used to analyze two independent groups, Kruskal-Wallis test was used to analyze multiple groups. Two-way repeated measures ANOVA was used to analyze different groups with two variables, including the sniffing on different odors, latency to the platform and total distance in open field test. All results showed were mean ± SEM, n represented the number of independent biological replicates and p value < 0.05 was considered significant. The statistical methods and statistical values of each result were presented in Supplementary Table 2.

## Author contributions

Jian-Ming Li and Tian-Ming Gao contributed to study concept; Jian-Ming Li, Tian-Ming Gao, Ying-Li, Jian-Ming Yang and Tong Shen contributed to study design; Ying-Li, Zheng-Yi Luo, Yu-Ying Hu, Yue-Wei Bi and Ming-An Liu contributed to data acquisition and analysis of behavioral and molecular studies; Zheng-Yi Luo and Ming-An Liu contributed to data acquisition and analysis of electrophysiological studies; Wen-Jun Zou and Yun-Long Song conducted the stereotactic surgery; Shu-Ji Li contributed to data acquisition of bacterial culture; Ying-Li and Lang-Huang conducted the HPLC; Shi-Li and Ai-Jun Zhou conducted the HE staining; Jian-Ming Li, Tian-Ming Gao, Ying-Li, Zheng-Yi Luo, Jian-Ming Yang and Yu-Ying Hu contributed to data analysis and interpretation; Jian-Ming Li, Tian-Ming Gao, Ying-Li and Yue-Wei Bi contributed to manuscript drafting; Jian-Ming Li contributed to funding obtaining and study supervision.

## Acknowledgements

This work was supported by National Key R&D Program of China [2017YFC1309000]; National Natural Science Foundation of China (Grant No. 81525020, 31570753, U1801282, 81901384); Guangzhou Science and Technology Plan Projects (Health Medical Collaborative Innovation Program of Guangzhou) [201803040019]; Natural Science Foundation of Jiangsu Province [BE2016666]; China Postdoctoral Science Foundation (Grant No. 2018M643319).

## Competion interests statement

The authors declare that there are no conflicts of interest.

## Data availability

The whole data in this work are available from the corresponding author upon reasonable request.

---

**Supplementary Figure 1.**
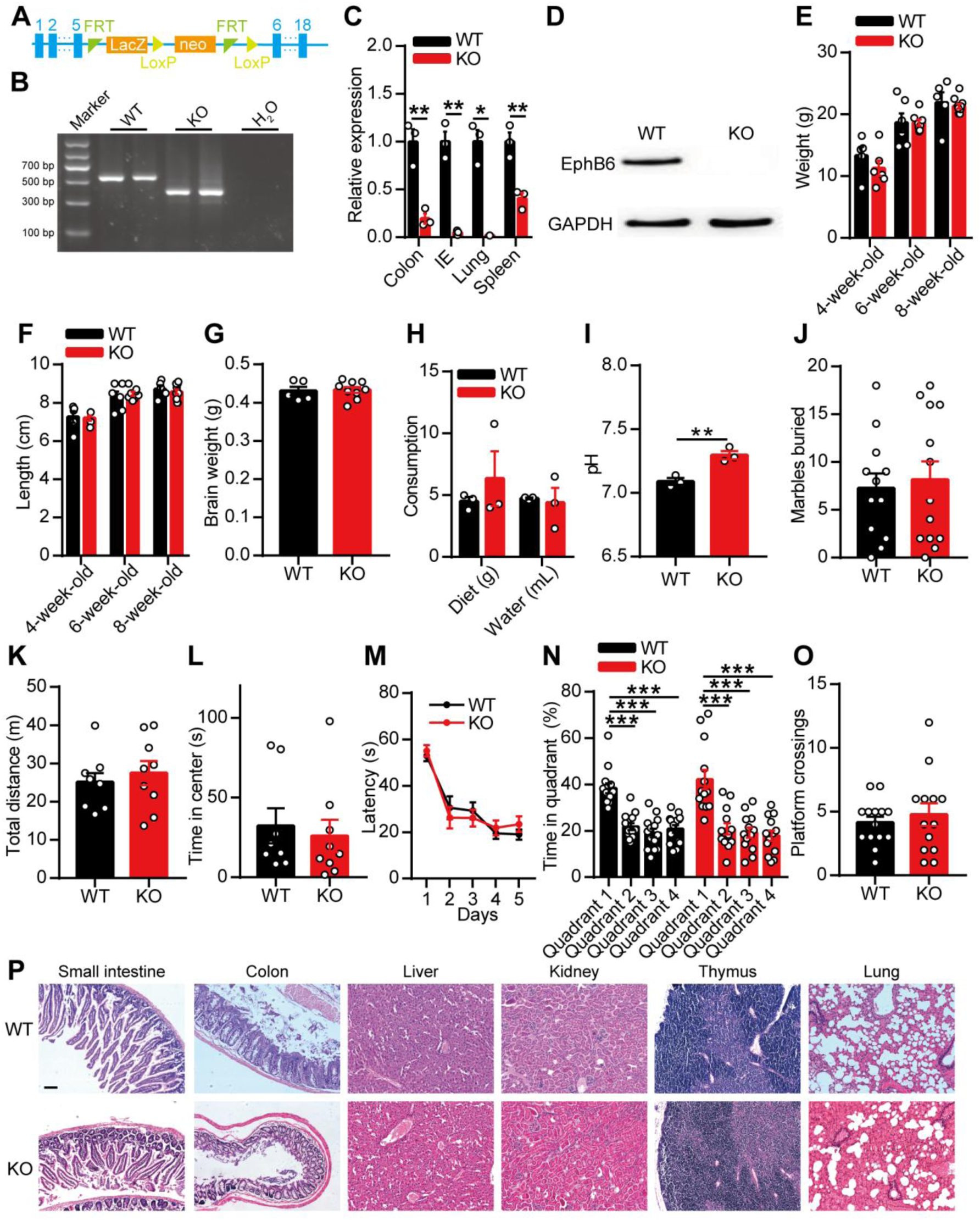
EphB6 knockout mice showed normal growth, motor behavior and spatial learning and memory. (A) Sequences inserted into the intron between exon 5 and exon 6 of EphB6. (B) Genotyping of WT and KO mice by agarose gel electrophoresis. The mutant band was 398 bp and the wild-type band was 542 bp. (C) The mRNA expression of EphB6 in different tissues of 8-week-old WT and KO mice. n = 3 mice for each group. (D) The protein expression of EphB6 in brain of 8-week-old WT and KO mice. (E-F) Similar body weight (E) and body length (F) between WT and KO mice. n = 5-10 mice for each group. (G) Similar brain weight between WT and KO mice. n = 5, 9 mice respectively. (H) Similar consumption of diet and water between WT and KO mice. n = 3 mice for each group. (I) The fecal pH of 8-week-old WT and KO mice. n = 3 mice for each group. (J) Marble burying test in 8-week-old WT and KO mice. n = 12, 13 mice respectively. (K-L) In open field test, total distances in 30 min (K) and time spent in center zone (L) between 8-week-old WT and KO mice were similar. n = 8, 9 mice respectively. (M-O) In morris water maze, 8-week-old KO mice spent same time to find the platform in acquisition test (M), spent same time in target quadrant (N) and had similar platform crossings (O) in probe test as WT mice. n = 14, 13 mice respectively. (P) The histological morphology of different organs of 8-weeks-old WT and KO mice by hematoxylin-eosin staining. Scale bar was 50 μm. Data shown are mean ± SEM. Two-tailed unpaired student’s *t* test (C, E-L, O), one-way ANOVA (N), two-way ANOVA (M). *, p < 0.05, **, p < 0.01, ***, p < 0.001. WT, EphB6^+/+^ mice; KO, EphB6^−/−^ mice, IE, intestinal epithelium. Statistical values are presented in Supplementary Table 2.

**Supplementary Figure 2.**
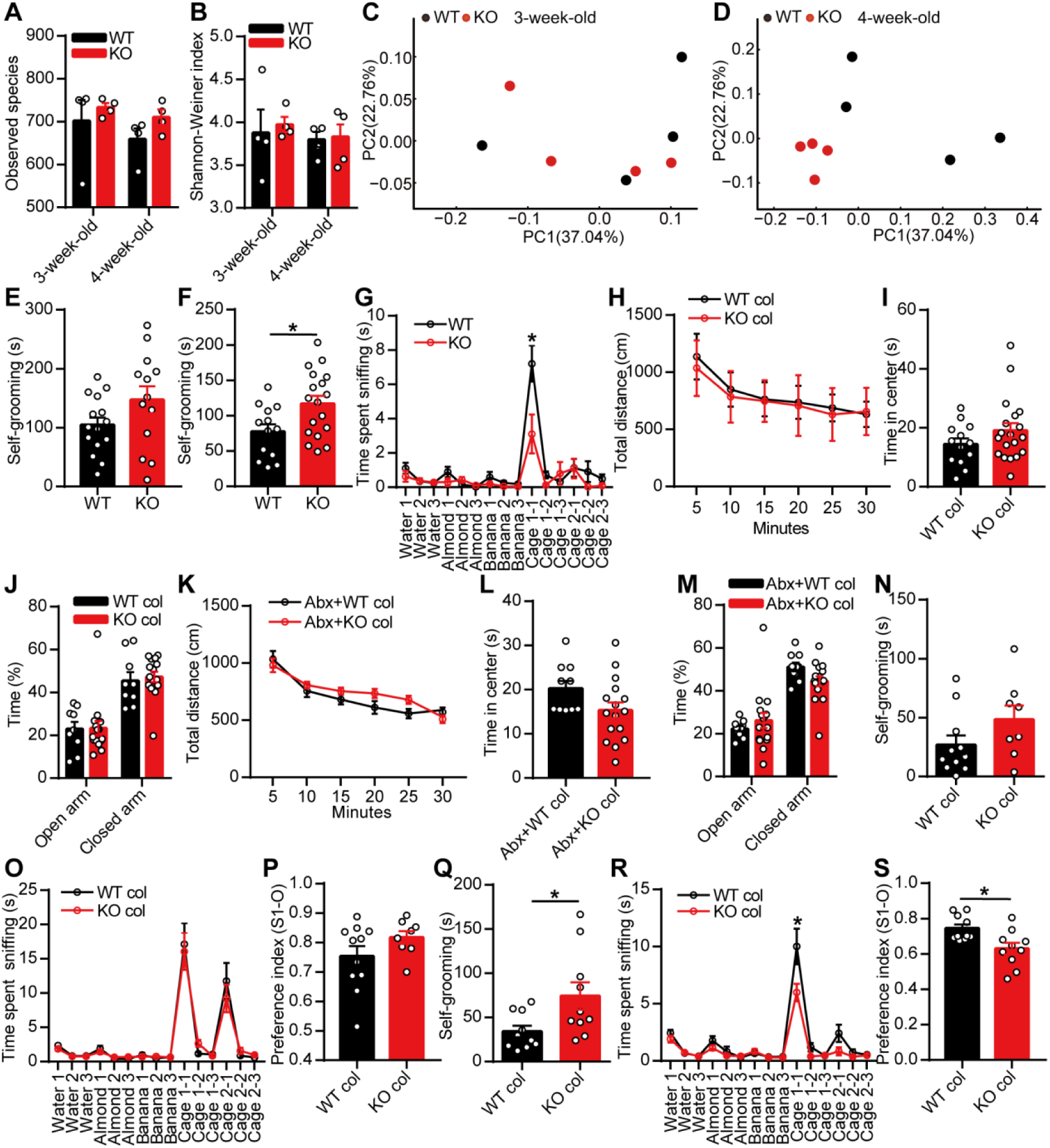
Fecal microbiota transplantation from 4-week-old EphB6 ablation mice induced autism-like behavior in SPF C57BL/6J mice. (A-D) 16S rDNA gene sequencing of gut microbiota of 3-4 weeks old WT and KO mice. The species richness (A) and diversity (B) of gut microbiota and the microbial composition (C-D) between the two groups. n = 4 mice for each group. (E) Self-grooming test in 3-week-old male WT and KO mice (21-23 days old). n = 15, 13 mice respectively. (F-G) Self-grooming test (b, n = 13, 18 mice respectively) and olfactory habituation/dishabituation test (c, n = 10, 12 mice respectively) in 4-week-old male WT and KO mice (27-29 days old). (H-J) Locomotor activities (H) and time in center zone (I) in open field test (H-I, n = 13, 19 mice respectively) and time spent in open arm and closed arm in elevated-plus-maze test (J, n = 9, 15 mice respectively) in 3-week-old SPF male C57BL/6J mice gavaged with fecal microbiota from 8-week-old male WT and KO mice. (K-M) Locomotor activities (K) and time in center zone (L) in open field test and time spent in open arm and closed arm in elevated-plus-maze test (M) in 3-week-old SPF male C57BL/6J mice gavaged with antibiotics and fecal microbiota from 8-week-old WT and KO mice. n = 10, 16 mice respectively. (N-P) Fecal microbiota from 3-week-old male WT and KO mice were gavaged to 3-week-old SPF male C57BL/6J mice for a week. After 3 weeks, self-grooming test (N), olfactory habituation/dishabituation test (O) and three-chambered social approach task (P) were conducted with an interval of at least 2 days. n = 11, 8 mice respectively. (Q-S) Fecal microbiota from 4-week-old male WT and KO mice were gavaged to 3-week-old SPF male C57BL/6J mice for a week. After 3 weeks, self-grooming test (Q), olfactory habituation/dishabituation test (R) and three-chambered social approach task (S) were conducted with an interval of at least 2 days. n = 10 mice for each group. Data shown are mean ± SEM. Two-tailed unpaired student’s *t* test (A-B, E-F, I-J, L-N, P-Q, S), two-way ANOVA (G-H, K, O, R), adonis analysis (C-D). *, p < 0.05. WT, EphB6^+/+^ mice; KO, EphB6^−/−^ mice; WT col or KO col, colonized with fecal microbiota from EphB6^+/+^ mice or EphB6^−/−^ mice; Abx, pre-treated with antibiotics (ampicillin, vancomycin, neomycin, metronidazole). Statistical values are presented in Supplementary Table 2.

**Supplementary Figure 3.**
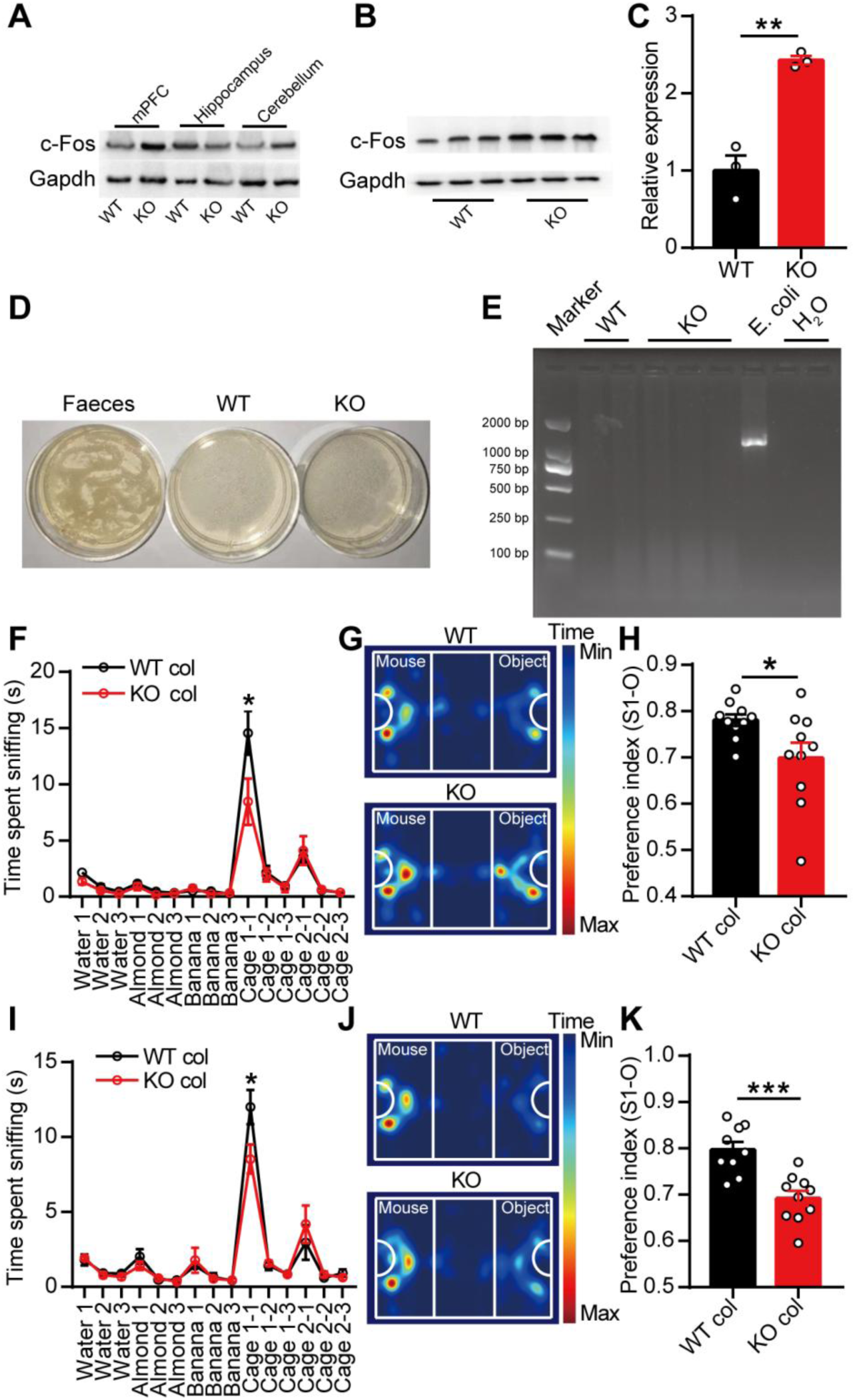
Both gut microbiota and their metabolites in EphB6 ablation mice induced social deficits in 6-week-old C57BL/6J mice. (A-C) The protein expression of c-Fos was detected in mPFC (B-C, n = 3 mice for each group), hippocampus and cerebellum of WT and KO mice 1 hr after being performed three-chambered social approach task. (D) Feces or PFC tissues of 8-week-old WT and KO mice were cultured in Luria-Bertani solid culture medium for 24 hr. n = 3,4 mice respectively. (E) DNA extracted from E. coli and PFC tissues of 8-week-old WT and KO mice were amplified by PCR using bacterial universal primers. n = 2, 3 mice respectively. (F-H) Fecal microbiota from 8-week-old male WT and KO mice was gavaged orally to 6-week-old SPF male C57BL/6J mice for a week. After a week, olfactory habituation/dishabituation test (F) and three-chambered social approach task (G-H) were conducted with an interval of at least 2 days. n = 10 mice for each group. (I-K) Fecal microbial metabolites from 8-week-old male WT and KO mice were gavaged orally to 6-week-old SPF male C57BL/6J mice for a week. After a week, olfactory habituation/dishabituation test (I) and three-chambered social approach task (J-K) were conducted with an interval of at least 2 days. n = 9, 10 mice respectively. Data shown are mean ± SEM. Two-tailed unpaired student’s *t* test (C, H, K), two-way repeated measures ANOVA (F, I). *, p < 0.05; **, p < 0.01; ***, p < 0.001. WT, EphB6^+/+^ mice; KO, EphB6^−/−^ mice; mPFC, middle prefrontal cortex; PCR, polymerase chain reaction; E. coli, *Escherichia coli*; WT col or KO col, colonized with fecal microbiota or metabolites from EphB6^+/+^ mice or EphB6^−/−^ mice. Statistical values are presented in Supplementary Table 2.

**Supplementary Figure 4.**
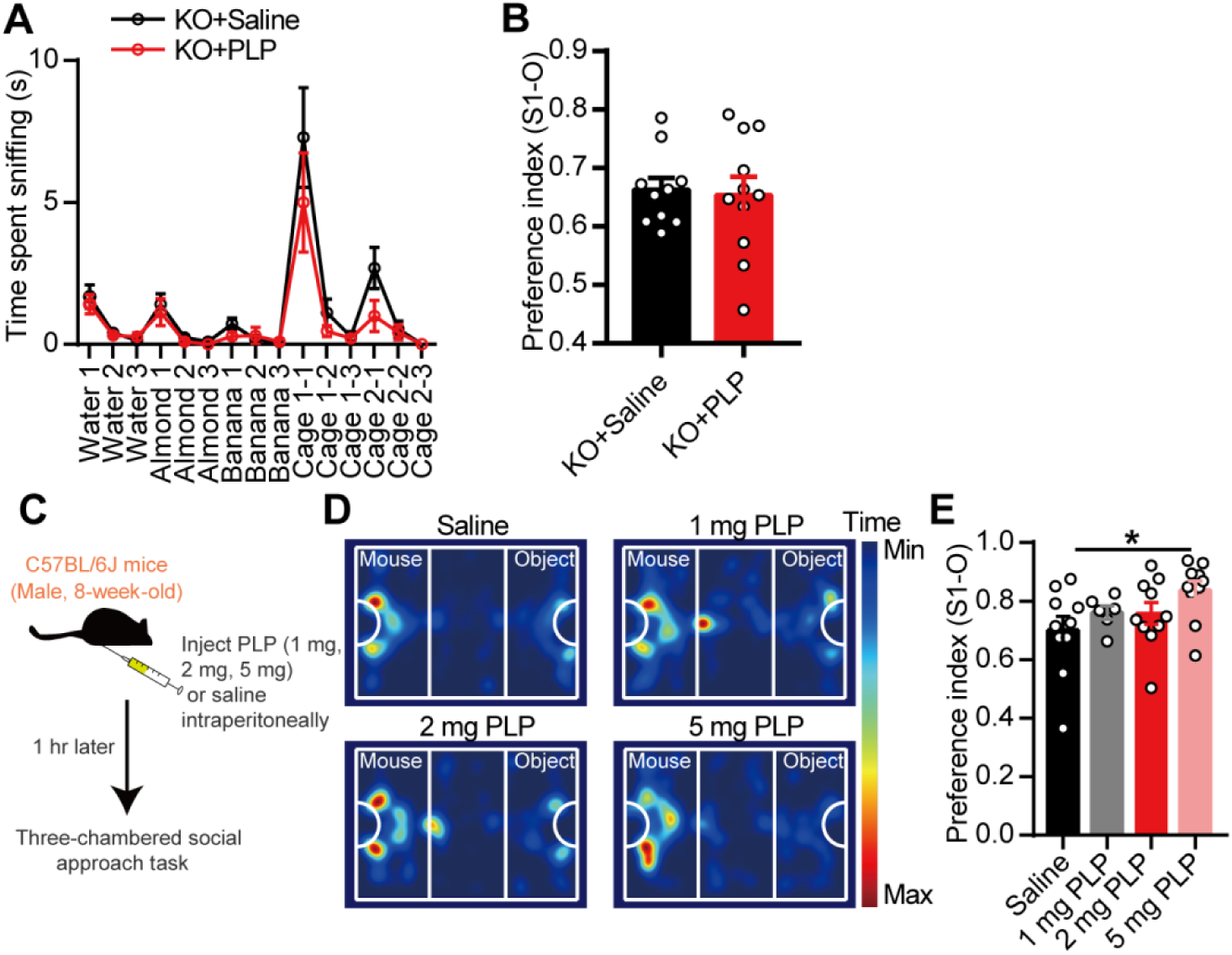
Vitamin B6 modulated social behavior in C57BL/6J mice. (A-B) 8-week-old male KO mice were gavaged with 1 mg PLP or saline. 2 hr later, olfactory habituation/dishabituation test (A, n = 12, 11 mice respectively) or three-chambered social approach task (B, n = 10, 11 mice respectively) was conducted. (C-E) Schematic of the injection of PLP (C). Saline or PLP (1 mg, 2 mg, 5mg per 0.2 mL) were injected intraperitoneally to 8-week-old male C57BL/6J mice. After 1 hr, three-chambered social approach task was conducted (D-E, n = 10, 6, 10, 10 mice respectively). Data shown are mean ± SEM. Two-tailed unpaired student’s *t* test (B), one-way ANOVA (E), two-way repeated measures ANOVA (A). *, p < 0.05. KO, EphB6^−/−^ mice; PLP, pyridoxal 5’-phosphate. Statistical values are presented in Supplementary Table 2.

**Supplementary Figure 5.**
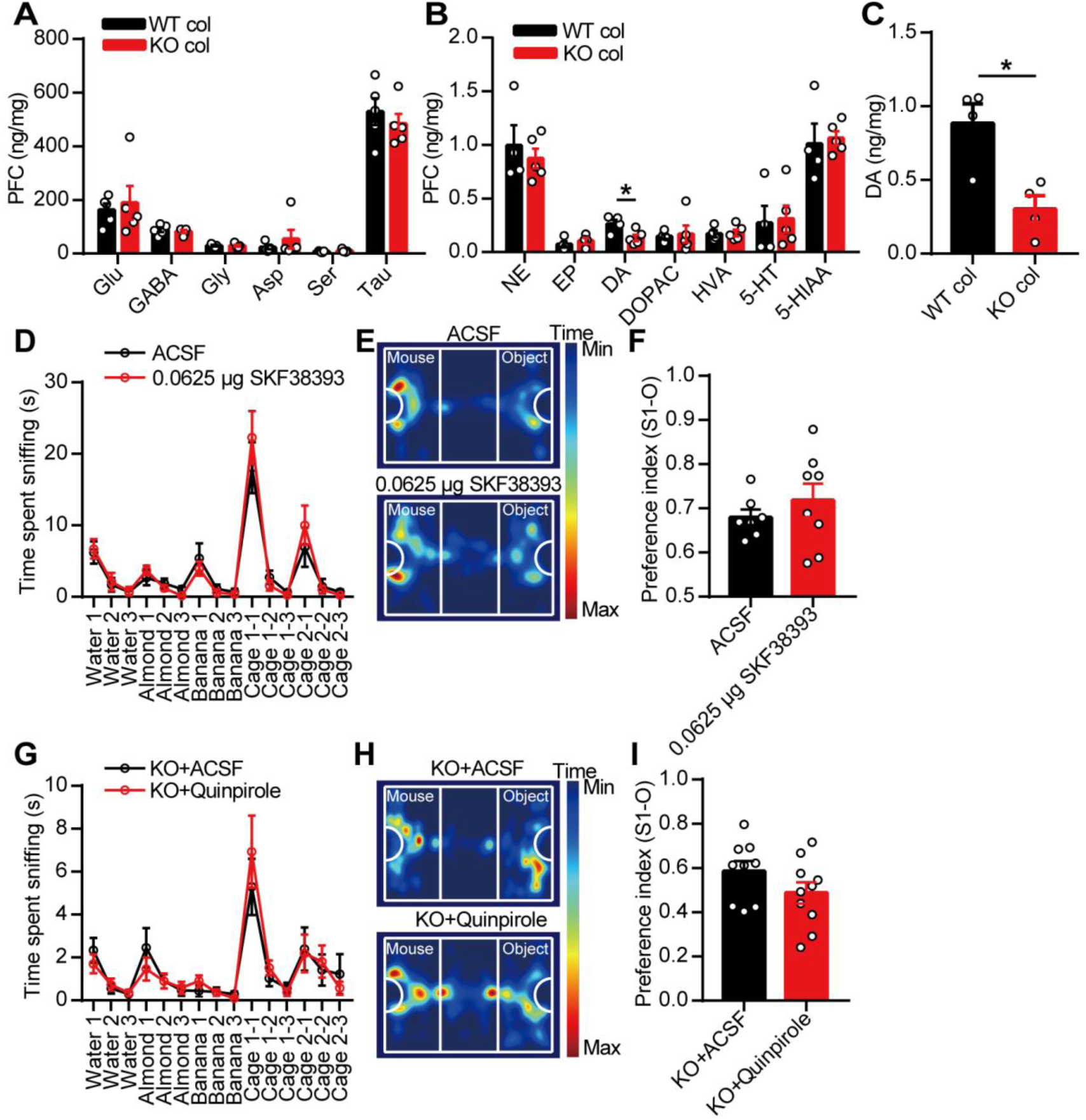
Dopamine was mediated by gut microbiota in PFC of mice. (A-B) 6-week-old SPF male C57BL/6J mice were gavaged with fecal microbiota from 8-week-old male WT or KO mice for a week, then a week later, the amino acid neurotransmitters (A) and monoamine neurotransmitters (B) in PFC of mice were detected by HPLC. n = 4-5 mice for each group. (C) Fecal microbiota of 8-week-old male WT and KO mice were gavaged to 3-week-old SPF male C57BL/6J mice for a week, then 3 weeks later, DA level in PFC of mice was detected. n = 4 mice for each group. (D-F) 8-week-old SPF male C57BL/6J mice injected with D1R agonist (SKF38393, 0.0625 μg/0.3 μL) in mPFC were performed with olfactory habituation/dishabituation test (D) and three-chambered social approach task (E-F) with an interval of a week. n = 7, 8 mice respectively. (G-I) Olfactory habituation/dishabituation test (G) and three-chambered social approach task (H-I) were conducted in 8-week-old male KO mice injected with D2R agonist (quinpirole, 1 μg/0.3 μL) or ACSF in mPFC with an interval of a week. n = 9, 10 mice respectively. Data shown are mean ± SEM. Two-tailed unpaired student’s *t* test (A-C, F, I), two-way repeated measures ANOVA (D, G). *, p < 0.05. WT col or KO col, colonized with fecal microbiota from EphB6^+/+^ or EphB6^−/−^ mice; PFC, prefrontal cortex; Glu, glutamic acid; GABA, gamma-aminobutyric acid; Gly, glycine; Asp, aspartic acid; Ser, serine; Tau, taurine; NE, norepinephrine; EP, epinephrine; DA, dopamine; 5-HT, 5-hydroxytryptamine; 5-HIAA, 5-hydroxyindoleacetic acid; DOPAC, dihydroxy-phenyl aceticacid; HVA, homovanillic acid; KO, EphB6^−/−^ mice; ACSF, artificial cerebrospinal fluid. Statistical values are presented in Supplementary Table 2.

**Supplementary Figure 6.**
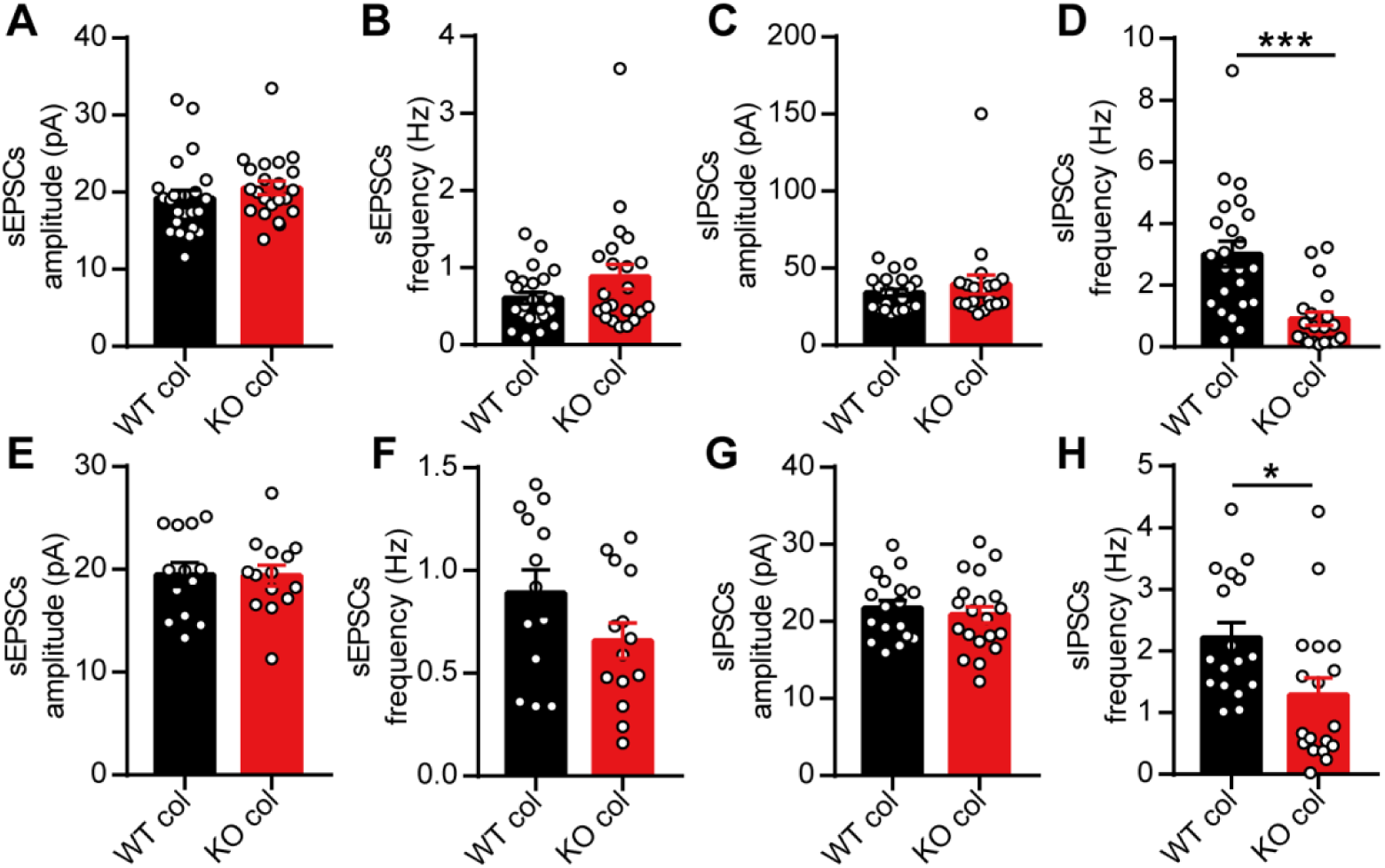
Gut microbiota modulated E/I balance in mPFC of C57BL/6J mice. (A-D) 3-week-old SPF C57BL/6J mice were gavaged with fecal microbiota from 8-week-old WT and KO mice for a week, 3 weeks later, we recorded sEPSCs and sIPSCs of pyramidal neurons in mPFC of mice. For sEPSCs, n = 23 and 22 cells from 4 mice respectively. For sIPSCs. n = 23 and 20 cells from 4 mice respectively. (E-H) 6-week-old SPF C57BL/6J mice were gavaged with fecal microbiota from 8-week-old WT and KO mice for a week, 1 week later, we recorded sEPSCs and sIPSCs of pyramidal neurons in mPFC of mice. For sEPSCs, n = 13 and 14 cells from 4 mice respectively. For sIPSCs. n = 17 and 18 cells from 4 mice respectively. Data shown are mean ± SEM. Two-tailed unpaired student’s *t* test (A-B, D-H), Mann-Whitney *U* test (C). *, p < 0.05; ***, p < 0.001. WT col or KO col, colocalized with fecal microbiota from EphB6^+/+^ or EphB6^−/−^ mice; mPFC, middle prefrontal cortex; sEPSCs, spontaneous excitatory postsynaptic currents; sIPSCs, spontaneous inhibitory postsynaptic currents. Statistical values are presented in Supplementary Table 2.

